# Neocortical astrocyte diversity stems from distinct developmental origins

**DOI:** 10.64898/2026.07.20.739583

**Authors:** Laura Dumas, Dorine Thobois, Jiafeng Zhou, Barbara Delaunay-Piednoir, Jason Durand, Inès Abdeddaim, Naim Khedair, Pauline Léger, Edson Rodrigues, Clémence Debacq, Emilie Pacary, David Ohayon, Hassan Boukhaddaoui, Martine Cohen-Salmon, Alexandre Pattyn, Riccardo Bocchi, Karine Loulier

## Abstract

Key regulators of neural network activity in multiple advanced cognitive processes and essential components of the blood-brain barrier, astrocytes constitute a highly heterogeneous population at the morphological, molecular, and functional levels. However, how this diversity arises during mammalian brain development remains poorly investigated. Here, using a combination of multicolour genetic fate mapping, single-cell transcriptomic analyses, multichannel large-volume imaging and detailed 3D cell morphology reconstructions, we uncover a discrete subpopulation of neocortical astrocytes generated from an early restricted embryonic domain located outside of the dorsal pallium. Besides their separate lineage from pyramidal neurons, these astrocytes exhibit a developmental trajectory that differs from astrocytes produced by dorsal cortical progenitors, including different migratory pathways, spatial distributions and morphology. Overall, our results reveal the diversity of embryonic sources responsible for neocortical astrocyte genesis and provide key insights into the unsuspected complex developmental processes that underlie cortical astrocyte heterogeneity.

## INTRODUCTION

For many years, complex brain functions such as cerebral plasticity or memory were reported to rely solely on neuronal network activity, whilst astrocytes were confined to supporting functions such as metabolic regulation and blood-brain barrier homeostasis. Key regulators of the formation and the regulation of neural circuits ^1–3^, astrocytes were nevertheless considered as mere facilitators rather than controllers of higher cognitive functions. This paradigm has now shifted thanks to recent studies demonstrating that astrocytes play a central role in many complex brain functions, forming local and long-range networks that connect multiple brain regions ^4^. Indeed, astrocytes are responsible for closing the critical period of visual plasticity ^5^, are essential modulators of emotional states under normal and chronic pain conditions ^6^ and constitute an active component of the engram, critical for memory storage and recall ^7^. In addition to these critical roles in close association with the neuronal circuitry, astrocytes are essential at the vascular interface, where they contribute to the maturation and the maintenance of the blood brain barrier ^8–10^, immune quiescence ^11,12^, and the regulation of neurovascular coupling ^13^. Supporting such numerous and diverse functions would require either a high level of subcellular computation within a homogeneous cell population or the existence of several specialised subpopulations, each dedicated to a specific function. Recent research verified both assumptions. Each astrocytic microdomains acts as a functional hub for multiple signal integration, receiving neuronal inputs from multiple synapses across distinct spatiotemporal scales at their perisynaptic (leaflets) processes ^14^, while also performing local translation at perivascular endfeet ^15^ simultaneously contacting blood vessels ^16^. Beyond their wide range of complex morphologies ^17–24^, cortical astrocytes display an extensive molecular heterogeneity, as revealed by single-cell and spatial transcriptomic analyses ^23,25–29^. Across central nervous system (CNS) regions, astrocyte functions rely heavily on both their molecular signature and their morphologies ^23^, both of which are shaped by local environmental cues provided by neighbouring neurons ^18^, the adjacent vasculature ^8^ and surrounding astrocytes ^17,30^. However, it is still unclear whether astrocyte diversity is already intrinsically predetermined ontogenetically or solely results from the myriad of factors to which astrocytes are exposed during CNS development. Unlike neurons, astrocytes display extensive plasticity both during healthy development ^20^ and in the mature brain in contexts of injury or neurodegenerative diseases ^23^. Until recently, cortical astrocyte production was assumed to rely mainly on the extensive proliferation of local astrocyte precursors ^31^ derived from regionally compartmentalised radial glia progenitors ^32^. Multicolour cell fate mapping strategies applied to track the progeny of multiple cortical progenitors have revealed considerable heterogeneity in the cellular composition, the size and spatial distribution of ontogenetically linked neocortical astrocytes, accompanied by significant postnatal maturation during the first month after birth, including increased morphological complexity and a larger territorial volume ^20^. These findings support the idea that astrocyte molecular identity and morphological features are acquired relatively late in development and are shaped by local environmental cues. However, a growing body of evidence suggests a more complex model for generating astrocyte diversity in the neocortex, whereby predetermined embryonic progenitors may contribute to specific astrocyte subpopulations. Twenty years ago, lineage tracing of the Dlx2-expressing cells arising from the ganglionic eminences revealed their contribution to astrocytes in the postnatal subventricular zone and cerebral cortex, without further characterising the astrocytes generated, nor their precise final location ^33–35^. These findings were later challenged by comprehensive genetic fate mapping analyses using multiple specific Cre mouse lines and precise viral targeting, demonstrating that neocortical astrocytes are generated exclusively by progenitors located in the dorsal pallium ^32^. More recently, studies combining targeted lineage tracing, genetic fate mapping, morphological analysis, and single-cell transcriptomics have provided evidence that temporally defined dorsal pallium progenitors were biased toward the generation of distinct neocortical astrocyte subpopulations ^36,37^. Astrocytes derived from late embryonic progenitors preferentially occupied upper cortical layers, whereas astrocytes generated by postnatal cortical progenitors were mainly located in the deep layers of the cerebral cortex. This suggests that the temporal patterning of cortical progenitors contributes to the laminar allocation of cortical protoplasmic astrocytes ^36^. This temporal bias in laminar and regional allocation has also been reported for other astrocyte subtypes, including fibrous and juxtavascular astrocytes ^37^. A comprehensive study of early cortical astrocyte heterogeneity has provided further evidence that two distinct dorsal pallial lineages contribute to molecularly and spatially defined astrocyte populations in the neocortex, using barcoding clonal lineage tracing together with single-cell and spatial transcriptomic analyses ^29^. Together, these studies support the idea that molecularly defined and temporally restricted dorsal pallial progenitors contribute to the spatial and molecular diversification of neocortical astrocytes. However, they do not yet capture the full extent of cortical astrocyte diversity, particularly given earlier evidence suggesting that non-dorsal pallial progenitors may also contribute to the generation of neocortical astrocytes. To reconcile these robust yet conflicting sets of evidence, it is essential to link the embryonic origins of specific neocortical astrocyte subpopulations to their developmental pathways, morphology, and final location.

Here, we discovered that a subset of cortical protoplasmic astrocytes (PrA) originates from embryonic progenitors located outside the dorsal pallium, supporting the hypothesis that developmental heterogeneity contributes to the diversity of cortical astrocytes. This new source was defined by combining multicolour genetic fate mapping, single-cell transcriptomic analyses, and large-volume imaging of cleared entire hemispheres to track astrogliogenesis from embryonic day (E)13.5 to postnatal day (P)60 during mouse forebrain development. Using multichannel multiphoton imaging ^38^ of CUBIC-cleared embryonic hemispheres from *Aldh1L1-CreER^T^*^2^ ^39^; *MAGIC Markers* floxed reporter mice^20,40^, we identified a discrete Aldh1L1-expressing embryonic domain near the pallium-subpallium boundary, which subsequently gives rise to astrocytes in the postnatal cerebral cortex. Neocortical astrocytes derived from these embryonic progenitors expressing Aldh1L1 at E13.5 disperse broadly across multiple areas of the postnatal cerebral cortex and preferentially settle in anterior and lateral neocortical regions, as well as within the deep cortical layers postnatally. Detailed three-dimensional reconstructions of individualised neocortical astrocyte arbors and volumes revealed that ventrally derived astrocytes (vA) display simpler morphologies than dorsal pallium-derived astrocytes (dA), despite occupying comparable territorial volumes during the first month after birth. Altogether, these results uncover unsuspected complex developmental mechanisms underlying the genesis of neocortical astrocyte heterogeneity relying on multiple embryonic sources, complex migratory pathways, and extended postnatal morphogenesis.

## RESULTS

### A subset of neocortical astrocytes originates from embryonic Aldh1L1-expressing cells

A comprehensive understanding of the neurodevelopmental origin of neocortical astrocytes requires a powerful lineage tracing strategy that enables the specific labelling of embryonic progenitors and tracking of their entire astrocyte progeny throughout development. In contrast to the traditional GFAP (glial fibrillary acidic protein), which labels only a subset of astrocytes in the adult mouse brain ^41^, aldehyde dehydrogenase 1 family member l1 (Aldh1L1) has emerged as a universal astrocyte marker through transcriptomic and immunohistochemical analyses ^42^. The *Aldh1L1-CreER^T^*^2^ BAC transgenic mouse line allows targeting of most astrocytes in the mature central nervous system, including protoplasmic astrocytes in the cortical grey matter ^39^. Although its effectiveness in tracking cortical astrocytes from early postnatal stages has been demonstrated ^43^, its usefulness for tracing astrocyte lineage in the mouse cerebral cortex from the embryonic stage has not yet been established, even though Aldh1L1 has been reported to be expressed prenatally in both mice and humans ^20,44^. To label cortical astrocytes from embryonic stages, *Aldh1L1-CreER^T^*^2^ mice ^39^ were crossed with the conditional multicolour *MAGIC Markers* reporter mice (MM ^20,40^) which enable the tracking of multiple individual cells based on the expression of distinct colour combinations (**Fig. 1; Supplementary Fig. 1**). Upon tamoxifen induction at P4, Cre recombination in *Aldh1L1-CreER^T^*^2^*;MM* pups triggers the expression of a combination of fluorescent proteins, resulting in multicolour labelling of astrocytes in the mouse cerebral cortex at P30 (**Fig. 1a-d**). Astrocyte identity was determined based on the characteristic bushy morphology of labelled cortical cells (**Fig. 1b-c**) and subsequently confirmed by Sox9 immunostaining (**Fig. 1d**), a nuclear transcription factor expressed by astrocytes across developmental stages ^45^. Consistent with previously reported labelling efficiencies in the adult-induced model ^39^, MM+Sox9+ astrocytes constituted 89.33 ± 1.03 % (mean ± SEM) of the total Sox9+ astrocyte population in the frontal and somatosensory neocortices at P30 (**Fig. 1e**). Within the MM+ population, 84.87 ± 1.54 % (mean ± SEM) were identified as Sox9+ cells, supporting the suitability of this strategy for targeting the astrocyte population of interest (**Fig. 1f**). No significant differences in astrocytic labelling ratio were observed between upper and deep cortical layers (**Fig. 1g**, upper layer ratio = 88.94 ± 1.19 % versus deep layer ratio = 89.79 ± 1.79 %) or across the medio-lateral axis (**Fig. 1h**, medial percentage of labelled astrocytes = 89.67 ± 1.04 % versus lateral percentage of labelled astrocytes = 88.00 ± 3.27 %).

**Fig. 1.**
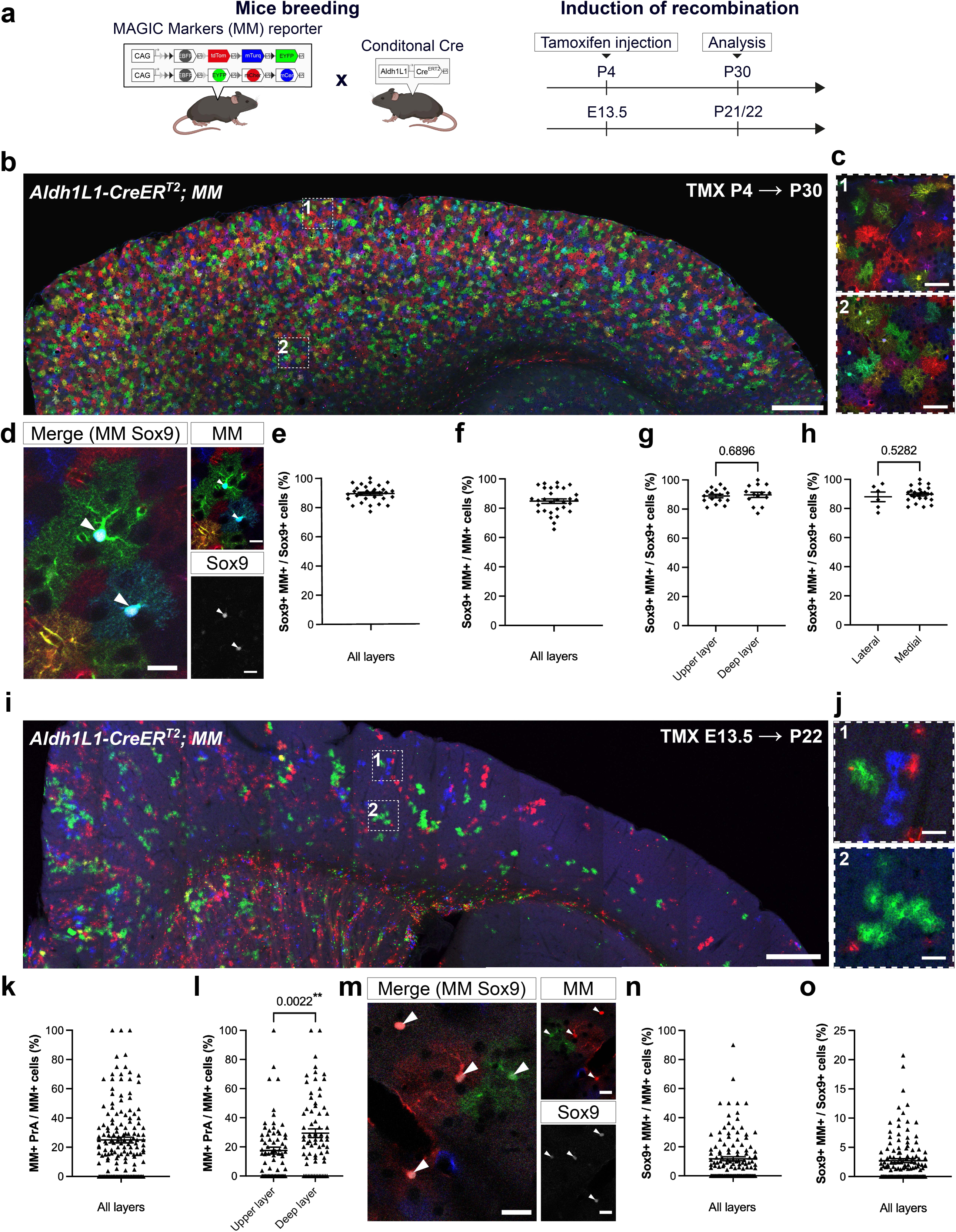
Lineage tracing using *Aldh1L1-CreER^T2^;MAGIC Markers* strategy reveals distinct astrocyte subpopulations in the postnatal cerebral cortex. **a** *MAGIC Marker (MM*) transgenic mice were crossed with *Aldh1L1-CreER^T2^*mice, and tamoxifen (TMX) was administered at either postnatal (P4) or embryonic (E13.5) stages to induce recombination. b *Aldh1L1-CreER^T2^;MM* neocortex at P30 after TMX-induced recombination at P4. **c** Close-up of the upper (**c1**) and lower (**c2**) layers from b. **d** Confocal image of MM-labelled astrocytes identified by co-labelling with Sox9 at P30 (TMX P4). **e** Quantification of MM^+^ astrocytes expressing Sox9 at P30 (TMX at P4). N = 3 independent animals, n = 30 areas from 9 sections. **f** Quantification of MM^+^ astrocytes among MM^+^ cells at P30 (TMX at P4). N = 3 independent animals, n = 30 areas from 9 sections. **g** Quantification of MM^+^ astrocytes between upper and deep layers at P30 (TMX at P4). Unpaired t-test, p = 0.6896, N = 3 independent animals, n = 16 and 14 areas from 9 sections. **h** Quantification of MM^+^ astrocytes between lateral and medial sections at P30 (TMX at P4). Unpaired t-test, p = 0.5282, N = 3 independent animals, n = 6 and 24 areas from 9 sections. **i** Confocal image of an *Aldh1L1-CreER^T2^;MM* mouse neocortex at P22 after embryonic induction of recombination with TMX administered at E13.5. **j** Close-up of upper (**j1**) and lower (**j2**) layers from i. **k** Quantification of MM^+^ protoplasmic astrocytes (PrA) among MM cells at P21/22 (TMX at E13.5). N = 15 independent animals, n = 152 areas from 36 sections. **l** Quantification of MM^+^ PrA between upper and deep layers at P21/22 (TMX at E13.5). Unpaired t-test, p = 0.0022, N = 14 independent animals, n = 72 areas per layer from 36 sections. **m** Confocal image of MM-labelled astrocytes identified by co-labelling with Sox9 at P21 (TMX E13.5). **n** Fraction of MM^+^ Sox9^+^ astrocytes among MM^+^ cells at P21/22 (TMX at E13.5). N = 11 independent animals, n = 128 areas per layer from 32 sections. **o** Percentage of labelled Sox9^+^ astrocytes among the total Sox9^+^ astrocyte population at P21/22 (TMX at E13.5). N = 11 independent animals, n = 128 areas per layer from 32 sections. Each count was conducted within regions of interest measuring 600 x 600 µm, distributed across upper and lower neocortical layers. All quantifications were done in the frontal and somatosensory cortical areas. Error bars represent means ± SEM. Scale bars: 500 (b, i); 50 (c, j) µm; 20 (d, m).

To characterise the early stages of cortical astrogliogenesis during embryogenesis, MM expression was induced by tamoxifen gavage of time-mated *Aldh1L1-CreER^T2^; MM* pregnant females at embryonic day (E)13.5, when Aldh1L1 begins to be expressed in the embryonic forebrain (**Supplementary Fig. 2a-d**). Brains were collected three weeks after birth to determine the fate of the progeny generated by E13.5 Aldh1L1-expressing cells (**Fig. 1a,i-j and Supplementary Fig. 2e-i**). Analysis of sagittal brain sections identified numerous MM-expressing cells exhibiting the characteristic bushy morphology of protoplasmic astrocytes (PrA) in multiple brain regions, including the cerebral cortex (**Fig. 1i-j**). In the frontal and somatosensory cortical areas, 24.92 ± 1.95 % (mean ± SEM) of MM-expressing cells were classified as PrA based on their distinctive bushy morphology (**Fig. 1k**), with a higher prevalence in the deep cortical layers (**Fig. 1l**). Among the cortical MM+ cells, 11.74 ± 1.38 % (mean ± SEM) expressed Sox9 (**Fig. 1m-n**) while the remaining MM+ neocortical cells were Sox9 negative and included interneurons and oligodendrocytes, but not pyramidal neurons (**Supplementary Fig. 1**). At P21/P22, MM-labelled cells accounted for an average of 2.72 ± 0.33 % (mean ± SEM), with marked variability ranging from 0 to 20.75%, of Sox9-positive cortical astrocytes in randomly sampled cortical areas and layers (**Fig. 1o**). These results indicate that E13.5 Aldh1L1-expressing embryonic progenitors generate a subset of protoplasmic astrocytes unevenly distributed in the postnatal cerebral cortex.

### Neocortical astrocytes originating from E13.5 Aldh1L1-expressing cells located outside the dorsal pallium enter the neocortex after E15.5 via tangential migration

To determine the location of the early progeny of E13.5 Aldh1L1-expressing cells, we first carried out lineage tracing experiments on E15.5 *Aldh1L1-CreER^T2^;MM* animals, which had been injected with tamoxifen at E13.5 (**Fig. 2a**). At this stage, MM+ cells were observed only in the ventral forebrain in the vicinity of ventricles (**Fig. 2a2**) and were absent from the dorsal forebrain (**Fig. 2a1**). In the absence of tamoxifen injection, no recombined MM+ cells were observed in E15.5 *Aldh1L1-CreER^T2^;MM* embryos (**Supplementary Fig. 3a**), demonstrating tight control of *MM* recombination by both Aldh1L1 promoter activity and the timing of recombination induction (E13.5). Of note, similar distributions of E13.5 Aldh1L1-derived progeny were detected using another well-described floxed single-colour reporter mouse line (**Supplementary Fig. 3b**). To track the location of the entire progeny of embryonic Aldh1L1-expressing cells without any loss of tissue material, we then performed whole-hemisphere CUBIC-based clearing^46^ of the brains of E15.5 *Aldh1L1-CreER^T2^;MM* embryos which received tamoxifen at E13.5 (**Fig. 2b**). This optical clearing approach was combined with multichannel multiphoton microscopy ^19,20,38^ to enable high-resolution volumetric tracking of MM+ cell dispersal within the intact hemisphere. Consistent with confocal analyses (**Fig. 2a**, **Fig. 2g**), MM+ cells at E15.5 were limited to ventral forebrain territories, with no MM-expressing cells in the dorsal forebrain throughout the entire hemisphere (**Fig. 2c-d**, **Supplementary movie 1**). At E15.5, MM+ cells were found mainly positioned close to the lateral ventricles in the ventricular zone extending from the pallial-subpallial boundary toward the medial ganglionic eminences (**Fig. 2c-d**, **Fig. 2h**). By E18.5, MM+ cells were found in greater quantities within the ganglionic eminences and have begun to enter the dorsal forebrain as early as E16.5 (**Fig. 2e-f**, **Fig. 2h-j; Supplementary movie 2**). In addition to the multiphoton imaging of CUBIC-cleared hemispheres, the daily tracking of MM+ cells from E15.5 to E18.5 using confocal imaging of brain coronal sections obtained from *Adh1L1CreER^T2^;MM* embryos that received tamoxifen at E13.5, revealed the progressive entry of MM+ cells into the dorsal forebrain via lateral and medial tangential migratory streams along the lateral ventricles (**Fig. 2g-j).** This combination of large-volume and high-resolution imaging using the CUBIC clearing technique, which preserves endogenous FP expression, enabled us to track the complexity of the migration pathways used by cells derived from Aldh1L1-expressing cells without any loss of information. Our work indicates that the progeny of E13.5 Aldh1L1-expressing progenitors are able to follow two distinct tangential routes to reach the neocortex: the lateral pathway used by interneurons ^47^ and the medial pathway taken by Dlx2+ cells ^33–35^.

**Fig. 2.**
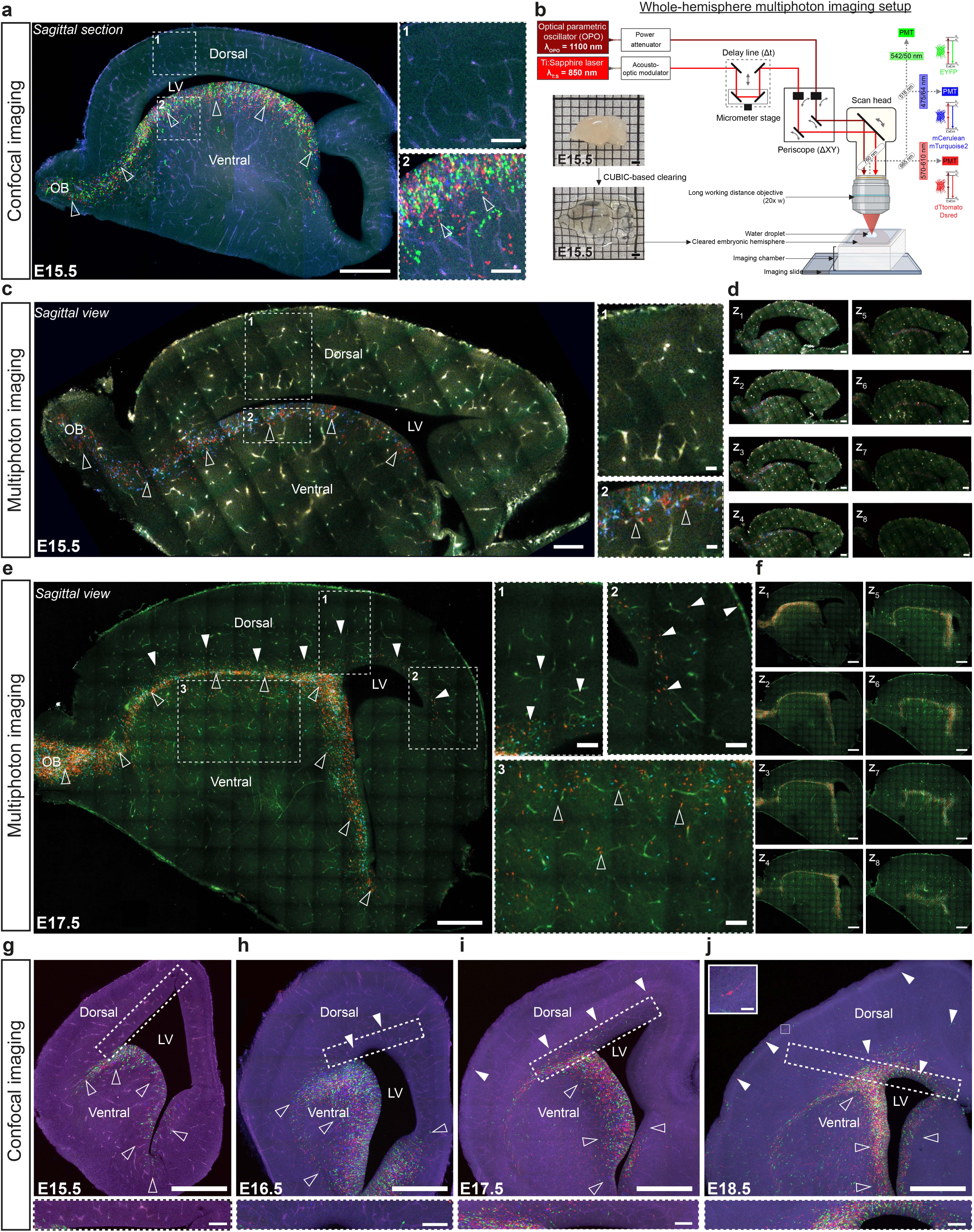
The progeny of E13.5 Aldh1L1-expressing cells colonises the mouse neocortex from the dLGE via tangential migratory pathways. **a** Confocal images of an *Aldh1L1-CreER^T2^;MM* sagittal brain section at E15.5 after TMX- induction of recombination at E13.5 (close-up of the dorsal (1) and ventral forebrain (2)). **b** CUBIC-based clearing of the entire embryonic hemisphere and multiphoton settings. **C** Multiphoton images of an *Aldh1L1-CreERT2; MM* mouse brain hemisphere at E15.5 (TMX at E13.5; detailed views of the dorsal (1) and ventral forebrain (2)). **d** Multiphoton serial images of the entire *Aldh1L1-CreERT2; MM* hemisphere at E15.5 (TMX at E13.5). **e** Two-photon serial images of an *Aldh1L1-CreERT2; MM* mouse brain hemisphere at E17.5 (TMX at E13.5; close-up of the dorsal (1,2) and ventral forebrain (3)). **f** Multiphoton serial images of the entire *Aldh1L1-CreER^T2^;MM* hemisphere at E17.5 (TMX at E13.5). **g** *Aldh1L1- CreER^T2^;MM* coronal section at E15.5 (TMX at E13.5) highlighting the ventricular zones without MM+ cells. **h** *Aldh1L1-CreER^T2^;MM* coronal section at E16.5 (TMX at E13.5), highlighting the ventricular zones with few MM+ cells. **i** *Aldh1L1-CreER^T2^;MM* coronal section at E17.5 (TMX at E13.5) showing a medial tangential migration along the ventricle and a lateral tangential pathway taken by MM+ cells. **j** *Aldh1L1-CreER^T2^;MM* coronal section at E18.5 (TMX at E13.5), highlighting the colonisation of the developing neocortex by numerous MM+ cells. Empty arrowheads denote subpallial localisation of MM+ cells, whereas full arrowheads signify pallial localisation of MM+ cells. The lateral tangential pathway is illustrated by a close-up of a migrating MM+ cell (upper close-up). LV = lateral ventricle, OB = olfactory bulb. Scale bar: 500 (a, c, d, e, f, g, h, i, j); 100 (a1, a2, c1, c2, e1, e2, e3, inset in g, inset in h, inset in i, lower inset in j); 20 (upper close-up in j) µm.

### Neocortical astrocytes derived from the embryonic ventral forebrain localise preferentially in frontal brain regions in the postnatal brain

Focusing on the neocortical astrocyte progeny derived from Aldh1L1-expressing progenitors located in the embryonic ventral forebrain, we next examined their rostro-caudal distribution within various brain regions at P7, P21, and P60 in *Aldh1L1-CreER^T2^;MM* animals after tamoxifen administration at E13.5 (**Fig. 3**). At P7 (**Fig. 3a-f**), the proportions of MM+ cells with PrA morphology over all MM+ cells assessed in rostral, intermediate, and caudal regions (**Fig. 3c**), were not significantly different across areas, even though a trend toward rostral bias was noticeable (13.55 ± 3.39 % MM+ PrA / MM+ cells in rostral versus 9.91 ± 2.42 % MM+ PrA / MM+ in intermediate and 4.03 ± 1.35 % MM+ PrA / MM+ cells in caudal region, **Fig. 3d**). Similarly, Sox9+ astrocytes originating from E13.5 Aldh1L1-expressing cells appeared to be distributed evenly along the rostro-caudal axis at P7, with comparable proportions over the total Sox9⁺ astrocytic population across the rostral (4.34 ± 1.17 % Sox9+ MM+/ Sox9+ cells), intermediate (3.42 ± 0.94 % Sox9+ MM+/ Sox9+ cells), and caudal (1.11 ± 0.48 % Sox9+ MM+/ Sox9+ cells) cortical regions. (**Fig. 3e**). However, these neocortical PrA derived from E13.5 Aldh1L1-expressing cells located in the embryonic ventral forebrain (vA) were found predominantly in lateral brain sections (11.34 ± 2.86 vA per mm^2^) compared to medial regions (**Fig. 3f**, 4.48 ± 0.35 vA per mm^2^). Collectively, these findings indicate that as early as P7, vA disseminate along the entire antero-posterior axis of the neocortex, with a preferential enrichment in lateral cortical regions and a non-significant tendency to favour the most rostral cortical regions. Two weeks later, at P21, the distribution pattern of MM+ vA had evolved to a graded arrangement along the rostro-caudal axis, with a significant preferential location in the most rostral area (**Fig. 3g-k**). Thus, anterior regions had a significantly higher proportion of MM+ cells with protoplasmic morphology (MM+ PrA) compared to posterior regions (**Fig. 3i**, 29.84 ± 2.99 % MM+ PrA / MM+ cells in rostral regions versus 8.85 ± 1.75% MM+ PrA / MM+ cells in caudal regions). Intermediate regions exhibited mixed values consistent with a gradient distribution (**Fig. 3i**, 20.00 ± 2.40 % MM+ PrA / MM+ cells, mean ± SEM). Likewise, Sox9+ vA also comprised a higher proportion of the total Sox9+ population in the rostral region (3.84 ± 0.57 % Sox9+ MM+/ Sox9+ cells) compared to the caudal region (**Fig. 3j** ; 0.63 ± 0.18 % Sox9+ MM+ / Sox9+ cells). Consistent with the lateral enrichment already observed at P7, vA were preferentially located in lateral regions (18.01 ± 1.86 vA per mm^2^) over medial brain regions (4.49 ± 0.85 vA per mm^2^) at P21 (**Fig. 3k**). These results indicate that vA display a non-random spatial distribution along both the rostro-caudal and medio-lateral cortical axis with a preferential location in the most rostral areas by P21. To determine whether this regionalisation of neocortical vA observed at P21 persists into maturity, their distribution was analysed at P60 (**Fig. 3l-p**). At this stage, MM+ vA remained significantly enriched in anterior (24.37 ± 5.46 % MM+ PrA / MM+ cells) areas compared to posterior regions (**Fig. 3n**; 7.04 ± 4.21 % MM+ PrA / MM+ cells). A similar distribution pattern was observed for Sox9+ vA (**Fig. 3o**), with a rostral enrichment (7.30 ± 1.80% Sox9+ MM+ /Sox9+ cells) that progressively decreased caudally (0.43 ± 0.24 % Sox9+ MM+/ Sox9+ cells). The density of vA also remained higher in lateral (2.11 ± 0.54 vA per mm^2^) compared to medial regions (1.44 ± 0.88 vA per mm^2^) at P60 (**Fig. 3p**). These results highlight a gradual postnatal colonisation of the neocortex by vA cells during the first three weeks, leading to their preferential localisation in rostral regions. This dynamic postnatal distribution of a PrA subpopulation (vA) derived from embryonic Aldh1L1-expressing cells in the ventral forebrain, reveals an unsuspected, protracted phase of cortical astrocyte dissemination during postnatal development.

**Fig. 3.**
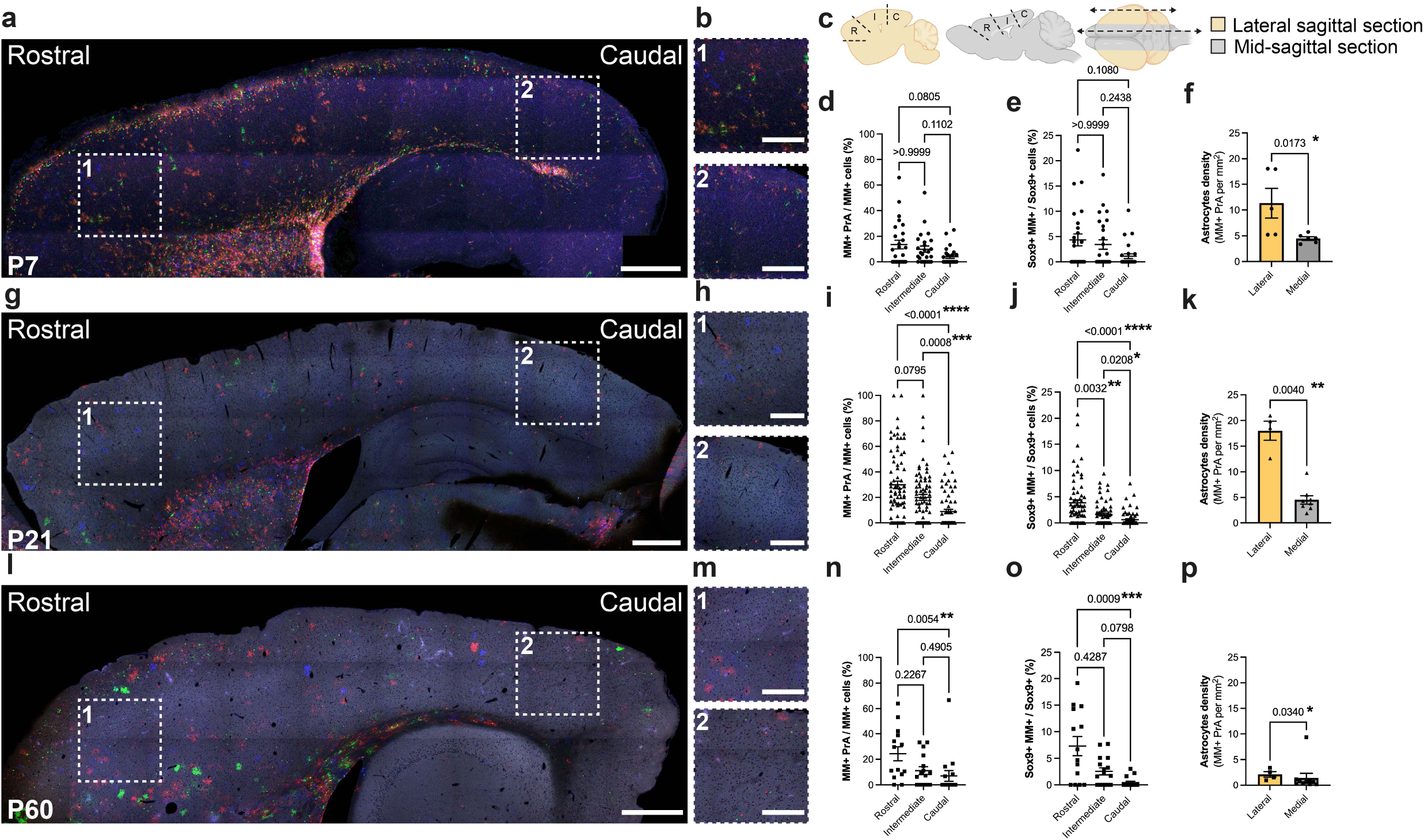
Spatial distribution of E13.5 Aldh1L1-expressing cell progeny across different regions of the postnatal neocortex. **a** *Aldh1L1-CreER^T2^;MM* sagittal section at P7 after E13.5 recombination. **b** Close-up of rostral (**b1**) and caudal (**b2**) regions from a. **c** Diagram of rostral (R), intermediate (I), and caudal (C) counting regions on lateral (yellow) and medial (grey) sagittal sections. **d** Percentage of MM^+^ protoplasmic astrocyte (PrA) among reporter^+^ cells across rostrocaudal cortical regions at P7 following E13.5 recombination. Kruskal-Wallis multiple comparison test, p-value for rostral versus caudal = 0.0805; p-value for rostral versus intermediate > 0.9999; p-value for intermediate versus caudal = 0.1102. N = 6 independent animals, n = 26 areas from 13 sections. **e** Fraction of Sox9^+^ MM^+^ astrocytes issued from E13.5 Aldh1L1-expressing cells within the Sox9⁺ population across cortical regions at P7 after E13.5 recombination. Kruskal-Wallis multiple comparison test, p-value for rostral versus caudal = 0.1080; p-value for rostral versus intermediate > 0.9999; p-value for intermediate versus caudal = 0.2438. N = 6 independent animals, n = 26 areas from 13 sections. **f** Quantification of MM⁺ PrA density along the medio-lateral axis at P7. Mann-Whitney test, p-value = 0.0173. N = 5 independent animals, n = 11 sections. **g** *Aldh1L1-CreER^T2^;MM* section at P21 after E13.5 recombination. **h** Close-up of rostral (**h1**) and caudal (**h2**) regions from g. **i** Percentage of MM^+^ PrA among MM^+^ cells across cortical regions at P21 after E13.5 recombination. Kruskal-Wallis multiple comparison test, p-value for rostral versus caudal < 0.0001; p-value for rostral versus intermediate = 0.0795; p-value for intermediate versus caudal = 0.0008. N = 14-15 independent animals, n = 72-76 areas from 37 sections. **j** Fraction of ventrally derived Sox9^+^ astrocytes among the Sox9⁺ cells across cortical regions at P21 after E13.5 recombination. Kruskal-Wallis multiple comparison test, p-value for rostral versus caudal < 0.0001; p-value for rostral versus intermediate = 0.0032; p-value for intermediate versus caudal = 0.0208. N = 11 independent animals, n = 64 areas from 32 sections. **k** Quantification of MM^+^ PrA density along the medio-lateral axis at P21. Mann-Whitney test, p-value = 0.0040. N = 6 independent animals, n = 18 sections. **l** *Aldh1L1-CreER^T2^; MM* section at P60 after E13.5 recombination. **m** Close-up of rostral (**m1**) and caudal (**m2**) regions from l. **n** Percentage of MM^+^ PrA among MM^+^ cells within cortical regions at P60 after E13.5 recombination. Kruskal- Wallis multiple comparison test, p-value for rostral versus caudal = 0.0054; p-value for rostral versus intermediate = 0.2267; p-value for intermediate versus caudal = 0.4905. N = 3 independent animals, n = 14-16 areas from 7 sections. **o** Fraction of Sox9^+^ MM^+^ astrocytes within the Sox9⁺ population across cortical regions at P60 after E13.5 recombination. Kruskal-Wallis multiple comparison test, p-value for rostral versus caudal = 0.0009; p-value for rostral versus intermediate = 0.4287; p-value for intermediate versus caudal = 0.0798. N = 3 independent animals, n = 14-16 areas from 7 sections. **p** Quantification of MM⁺ protoplasmic astrocyte density along the medio-lateral axis at P60. Mann-Whitney test, p- value = 0.0340. N = 5 independent animals, n = 4-10 sections. Error bars represent means ± SEM. Scale bars: 500 (**a**, **g**, **l**); 250 (**b**, **h**, **m**) µm.

### Developmental origin defines distinct morphological features of neocortical astrocytes

As neocortical astrocytes derived from Aldh1L1-expressing embryonic cells display a differential spatial distribution across distinct cortical regions in the postnatal brain, we investigated whether vA may exhibit distinct morphologies compared to neocortical astrocytes derived from dorsal pallium progenitors (dA). The morphological features, including 3D volume and arborisation complexity of the two subpopulations at the single-cell level, were assessed using MM expression. As previously described, dA were labelled by in utero electroporation of MM transgenes in E15.5 cortical progenitors located in the dorsal pallium ^20,24,40,48^, whereas vA were marked using tamoxifen induction of *Aldh1L1-CreER^T2^;MM* embryos at E13.5 (**Fig. 1-3**). Brains were collected at P7 (**Fig. 4a-e**), P21 (**Fig. 4f-j**), and P60 (**Fig. 4k-o**). Confocal microscopy was used to acquire high-resolution 3D z-stacks that captured the complete three-dimensional morphology of individual astrocytes in the frontal and somatosensory cortical regions over a volume of several tens of cubic micrometres, with no significant morphological differences in MM+ PrA analysed in these two cortical areas. The expression of MM fluorescent proteins allowed for reliable individualisation and tracking of astrocyte processes and delineation of their full territorial domains using dedicated software (IMARIS for territorial volume and Vaa3D for arbor complexity). Up to P21, dA and vA occupied comparable territorial volume (at P7: dA 9841 ± 1031 µm³ vs vA 17131 ± 3971 µm³, **Fig. 4b**; at P21: dA 52346 ± 3545 µm^3^ vs vA 53172 ± 3892 µm^3^, **Fig. 4g**). At P60, however, dA territories were larger than those of vA (dA 57117 ± 2340 µm³ vs vA 45055 ± 3271 µm³, **Fig. 4l**), revealing a late divergence in territorial volume between the two populations. At P7, vA have a morphology comparable to dA, characterised by similar number of nodes (vA 3905 nodes ± 774 vs dA 3973 ± 492; **Fig. 4c**), branches (vA 2077 branches ± 439 vs dA 2664 ± 460; **Fig. 4d**), and terminal tips (vA 1122 tips ± 242 vs dA = 1438 ± 254; **Fig. 4e**). At P21, even though both vA and dA volume and arbour complexity increase as expected from previous studies^17,20^, vA exhibit a significantly simpler morphology compared with dA with a reduced number of nodes (vA 9313 ± 1423 vs dA 13628 ± 895; **Fig. 4h**), branches (vA 5338 ± 1060 vs dA 9633 ± 660; **Fig. 4i**), and tips (vA 2911 ± 601 vs dA 5182 ± 363; **Fig. 4j**). To determine whether the reduced arbour complexity of vA up to P21 reflected a transient developmental delay that might resolve with further maturation, we analysed vA morphological features at P60. At this stage, dA exhibit a larger territorial volume than vA (dA = 57117 ± 2340 µm³ versus vA = 45055 ± 3271 µm³; **Fig. 4l)** but no more significant differences in arborisation metrics (**Fig. 4m-o**), indicating a convergence of arbour features at this stage. However, deeper analysis of vA and dA morphological developmental trajectories focusing on their late morphogenesis, revealed a significant reduction in dA arborisation complexity between P21 and P60 (**Supplementary Fig. 4a-d**) whereas vA morphological features do not change markedly during this period. With regard to territorial volumes, dA exhibit a smaller volume in the deep layers (44637 ± 1977 µm³) than in the upper layers (61598 ± 4900 µm³) at P21, which is no longer visible at P60, whilst vA, which show no difference in domain size between layers at P21, exhibit a larger volume when they are located in the deep layers at P60 (upper dA = 57994 ± 2308 µm^3^ vs deep dA = 56239 ± 4233 µm^3^; **Supplementary Fig. 4e-f**). At P21, vA displayed no difference in domain size between layers (upper vA = 51478 ± 4266 µm^3^ vs depp vA = 54019 ± 5612 µm^3^), but demonstrated a larger volume when positioned within the deep layers at P60 (deep vA = 49691 ± 3270 µm^3^ vs upper vA = 35784 ± 3239 µm^3^; **Supplementary Fig. 4e-f**). These results indicate that dA and vA undergo distinct developmental trajectories regarding their morphological maturation at postnatal stages. They also reveal an unexpected late phase of maturation for dA after P21 that relies on the simplification of their morphology, likely in line with their functions at synaptic or vascular interfaces.

**Fig. 4.**
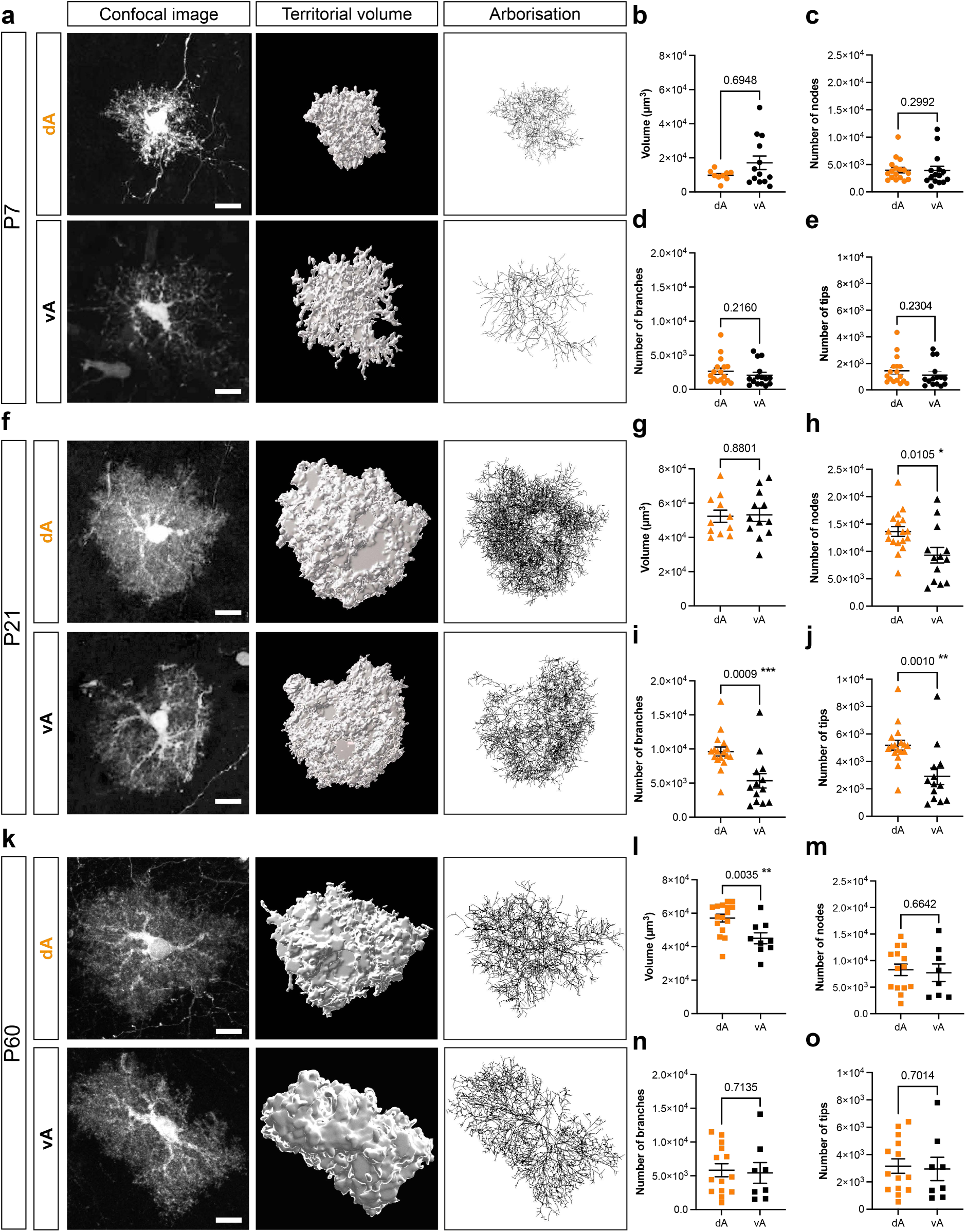
Ontogenetically distinct neocortical astrocytes display different morphological features and developmental trajectories. **a** 3D reconstructions from confocal images illustrating the arborisation and territory of astrocytes originating from either the dorsal pallium (dA) or the ventral Aldh1L1-expressing embryonic ventral domain (vA) at P7. **b** Astrocyte territorial volume between dA and vA at P7. Mann-Whitney test, p = 0.6948; N = 5 (dA) and 3 (vA) independent animals, n = 9 (dA) and 13 (vA) astrocytes. **c, d, e** Number of nodes, branches, or tips, respectively, between dA and vA at P7. Mann-Whitney test, p = 0.2992 (nodes); p = 0.2160 (branches), p = 0.2304 (tips); N = 5 (dA) and 5 (vA) independent animals, n = 17 (dA) or 15 (vA) astrocytes. **f** 3D reconstructions of arborisation and territory of dA and vA at P21. **g** Astrocyte territorial volume between dA and vA at P21. Mann-Whitney test, p = 0.8801, N = 4 (dA) and 8 (vA) independent animals, n = 11 (dA) and 12 (vA) astrocytes. **h, i, j** Number of nodes, branches and tips, respectively, between dA and vA at P21. Mann-Whitney test, p = 0.0105 (nodes); p = 0.0009 (branches); p = 0.0010 (tips); N = 4 (dA) and 7 (vA) independent animals, n = 17 (dA) and 13 (vA) astrocytes. **k** 3D reconstructions of arborisation and territory of dA and vA at P60. **l** Astrocyte territorial volume between dA and vA at P60. Mann-Whitney test, p = 0.0035, N = 4 (dA) and 5 (vA) independent animals, n = 16 (dA) and 9 (vA) astrocytes. **m, n, o** Number of nodes, branches and tips, respectively, between dA and vA at P60. Mann- Whitney test, p = 0.6642 (nodes); p = 0.7135 (branches); p = 0.7014 (tips); N = 3 (dA) and 4 (vA) independent animals, n = 14 (dA) and 8 (vA) astrocytes. All quantification were done in the frontal and somatosensory cortical areas. Error bars represent means ± SEM. Scale bars: 10 µm.

### Despite differential spatial distribution and morphological differences, ventrally-derived neocortical astrocytes share similar features with dorsally-derived astrocytes

As vA and dA display distinct developmental trajectories, we wondered whether vA may also exhibit differential maturation at the vascular interface. We therefore examined their integration into the vascular compartment, using blood-vessel contact and perivascular aquaporin polarization as readouts of astrocyte-vascular interface maturation in the postnatal neocortex. At P7, MM-labelled vA (MM-vA), whose identity was confirmed by Sox9 co-expression, were observed in close apposition to CD31-positive blood vessels, although some neighbouring Sox9+ MM-labelled cells lacked overt vascular contact (**Fig. 5a**). vA were associated with small CD31+ vessels but were not observed contacting large SMA+ vessels (data not shown). By P21/22, MM-vA maintained extensive associations with the vascular network, as demonstrated by three-dimensional reconstruction of individual astrocytes and CD31-positive vessels (**Fig. 5b**). In line with the development of the perivascular astrocytic interface, AQP4 was detected in MM-vA at P7 (**Fig. 5c**) and became polarised at vascular contact sites by P21/22 (**Fig. 5d**). Thus, despite their distinct developmental trajectories, vA exhibit a similar maturation at the gliovascular interface, including vascular contacts and perivascular AQP4 polarisation, on a timeline comparable to dA vascular maturation typically observed in the postnatal cortex ^49^. We then assessed the molecular identity of vA at P21 using canonical astrocytic markers, including Aldh1L1, GFAP or Olig2. vA displayed heterogeneous expression of astrocytic markers, with a limited fraction of MM⁺ PrA expressing Aldh1L1 (89.16 ± 3.65%; **Fig. 5e**), GFAP (39.28 ± 7.64%; **Fig. 5f**), or Olig2 (42.55 ± 6.35%; **Fig. 5h**). Taken together, these data indicate that, despite their distinct spatial distribution and morphological maturation, vA acquire key perivascular astrocytic features at a developmental pace similar to dA, while exhibiting molecular heterogeneity at the population level in the postnatal neocortex.

**Fig. 5.**
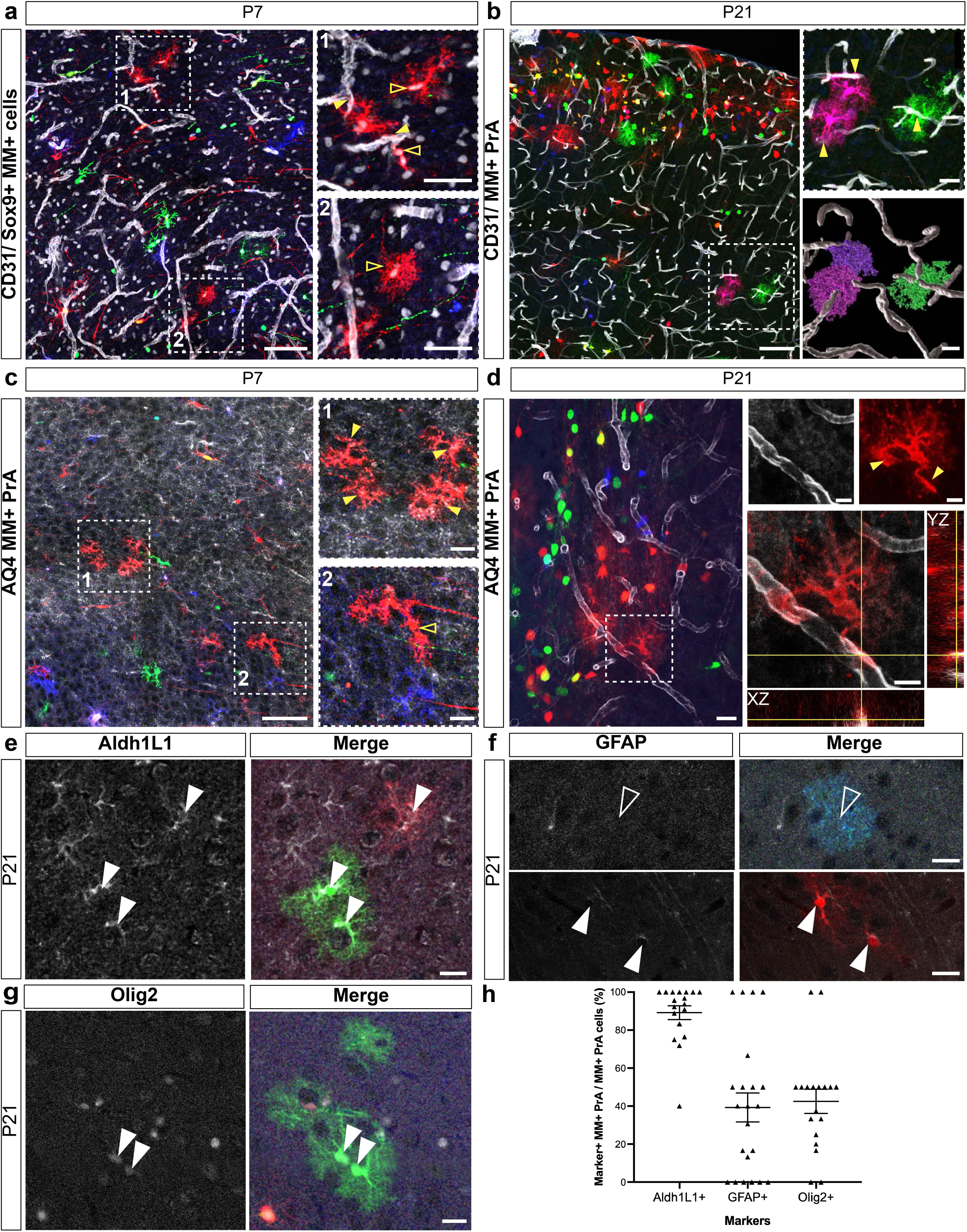
Aldh1L1-derived neocortical astrocytes acquire timely perivascular features and display molecular heterogeneity. **a** Confocal image at P7 showing MM-labelled astrocytes issued from the Aldh1L1-expressing embryonic domain (vA), identified by nuclear Sox9 co-labelling (grey), together with CD31- labelled blood vessels (grey). Arrowheads indicate Sox9^+^ vA contacting CD31^+^ blood vessels, whereas stars mark MM^+^ Sox9^+^ cells lacking vascular contact. N = 4 mice; 18- 84 MM^+^ cells per mouse. **b** Confocal images at P21) illustrating vA. Arrowheads denote vascular contact sites (CD31, grey). At the bottom, a 3D rendering of three reconstructed astrocytes and the vascular network is provided. N = 3 mice. **c** Confocal image at P7 depicts vA expressing AQP4 (grey). Arrowheads denote AQP4^+^ vA contacting blood vessels, whereas cells marked by yellow stars indicate cells lacking vascular contact. **d** Confocal image at P21 showing vA with AQP4 polarized at the vascular interface (grey). Arrowheads indicate perivascular AQP4^+^ astrocytes. Orthogonal (YZ and XZ) views illustrating vA and AQP4 co- localization (red and grey, respectively). **e, f, g, h** Molecular characterization of MM-vA at P21. Confocal images showing vA expressing Aldh1L1 (**e**, grey), not expressing GFAP (**f** top, grey), expressing GFAP (**f** bottom, grey) or expressing Olig2 (**g**, grey). **h**, Quantification of the proportion of MM^+^ PrA deriving from the ventral forebrain expressing each marker within the MM^+^ PrA population. Error bars represent means ± SEM, N = 4 (Aldh1L1, GFAP, Olig2) independent animals, n = 18 (Aldh1L1), 22 (GFAP), 18 (Olig2) areas from 9-11 sections. Boxed regions are shown at higher magnification. Vascular interface quantifications were performed in the rostral cortical region, whereas molecular marker quantifications were performed across the entire cortex. Aldh1L1: Aldehyde dehydrogenase 1 family member L1, AQP4: Aquaporin 4, GFAP: Glial Fibrillary Acidic Protein, Olig2: Oligodendrocyte transcription factor 2. AQP4: Aquaporin 4, GFAP: Glial Fibrillary Acidic Protein. Scale bars: 20 µm (a, b, c, d, e, f, g), 10 µm (a1, a2, insets in d), 5 µm (insets in b).

### Embryonic Aldh1L1-expressing cells are found in a spatially and temporally regulated domain that straddles ventral pallium and subpallium territories

We showed that a subpopulation of neocortical astrocytes displaying distinct spatial distribution, morphologies and developmental trajectories was derived from embryonic Aldh1L1-expressing cells located in the ventral forebrain, suggesting an ontogenetic origin that differs from the one usually described for neocortical astrocytes, namely radial glia cells located in the dorsal pallium ^20,21,32,50^. To better characterise the embryonic Aldh1L1-expressing cells, we first determined the *Aldh1L1* gene expression pattern by in situ hybridisation (ISH) from E13.5 to E18.5. While no expression of Aldh1L1 is reported at E12.5 (**Supplementary Fig. 2a**), at E13.5, *Aldh1L1* mRNA is present in a restricted domain located in the dorsal part of the lateral ganglionic eminence (dLGE) and on the ventral side of the pallial-subpallial boundary (PSB), which separates the developing neocortex from the ganglionic eminences (GE, **Fig. 6a-b**). By E15.5, the Aldh1L1-expressing domain expanded ventrally, lining the lateral ventricle, towards the medial ganglionic eminences. By E18.5, the Aldh1L1 domain surrounded the entire lateral ventricles, including the dorsal and remaining medial wall of the ventricle (**Fig. 6b**). The dynamic Aldh1L1 mRNA expression pattern is mirrored by Aldh1L1 protein expression revealed by immunostaining at all embryonic stages (**Supplementary Fig. 2a-d**) and confirmed using *Aldh1L1-eGFP* embryos ^51^ (**Supplementary Fig. 5a-c**). Although anatomical landmarks provide a framework for defining embryonic domains, these territories are more precisely delineated by a combination of transcription factors (TF; **Fig. 6a,c**). To determine the molecular identity of Aldh1L1-expressing cells at E13.5 and their potential correspondence to known domains, we carried out sequential double ISH. We used probes specific to TF expressed in the dorsal, lateral, and medial pallium (*Emx1*, *Neurog2*, *Pax6*, *Tfap2c*), ventral pallium (*Sfrp2*), and subpallium (*Ascl1*, *Dlx2*). *Adh1L1* expression partially overlapped with several TFs: *Neurog2*, *Pax6*, *Tfap2c*, *Sfrp2*, *Ascl1*, and *Dlx2* (arrowheads, **Fig. 6c**). However, no overlap was observed between *Aldh1L1* and *Emx1* (**Fig. 6c**) mRNA expression patterns, suggesting an exclusion of the Aldh1L1-expressing domain from the Emx1-defined dorsal pallium at this stage. These results indicate that the Aldh1L1-expressing domain at E13.5 spans a molecularly complex region located in the dorsal part of the lateral ganglionic eminence encompassing part of the ventral pallium (*Pax6*+, *Sfrp2+, Emx1-)* and part of the subpallium (*Ascl1+*, *Dlx2+)*, but distinct from the dorsal pallium (**Fig. 6d**). We then investigated the molecular identity of E13.5 Aldh1L1-expressing cells at the protein level (**Fig. 6e-g)**. Mirroring Aldh1L1 mRNA expression, Aldh1L1 protein expression starts around E13.5 in the dLGE and extends ventrally toward the medial ganglionic eminences for the next two days (**Supplementary Fig. 2a-d).** This expression pattern is consistent with our lineage tracing experiments relying on tamoxifen administration using either single (at E12, E13 or E15) or cumulative injections (E13+E14+E15), which show that whilst no astrocytes are found in the neocortex of *Aldh1L1CreERT2;MM* injected at E12, a very large number of MM+ cortical astrocytes are found following cumulative injections, along with a greater number of interneurons (**Supplementary Fig. 2e-i)**. Co-immunostaining for Aldh1L1 and Pax6 proteins in E13.5 mouse brains reveals two levels of Pax6 expression in the dLGE, Pax6^high^ (**Fig. 6e1**) and Pax6^low^ (**Fig. 6e2**), with Aldh1L1 expressed mostly in the Pax6^low^ population : 89.74 ± 2.16% of Aldh1L1-positive cells expressed low levels of Pax6, whereas only 5.16 ± 2.09% expressed high levels (**Fig. 6e3**). Interestingly, a small fraction of the Aldh1L1+ population expressed the TF Dlx2 (4.06 ± 1.17%; **Fig. 6f-g**), confirming that a subset of the Aldh1L1+ embryonic cells have a subpallial identity. Because Dlx2 is re-expressed in the postnatal brain and no conditional Cre lineage tracing mouse line is available, the *Ascl1-CreER^T2^* mice ^52^ were used to track the fate of E13.5 Ascl1+ cells located in GE using our multicolour MM reporter mice (**Supplementary Fig. 6**). MM reporter mice were crossed with *Ascl1-CreER^T2^* mice, with tamoxifen administered at E13.5 and analyses performed at E15.5, E18.5, and P8 (**Supplementary Fig. 6**). Aldh1L1- and Ascl1-derived cells showed distinct distribution patterns at all stages. At E15.5, Aldh1L1-derived MM+ cells were located close to the lateral wall of the lateral ventricles (**Supplementary Fig. 6a**), while Ascl1-derived cells were broadly distributed in the developing striatum (**Supplementary Fig. 6b**). By E18.5, Ascl1-derived cells had dispersed across presumptive striatal and septal regions (**Supplementary Fig. 6d**), while Aldh1L1-derived cells displaying a radial glia morphology, remained enriched in the ventricular zone and expanded across medial ganglionic eminences on one side and towards the dorsal wall of the lateral ventricles (**Supplementary Fig. 6c**). At P7-P8, Aldh1L1-derived cells populated the cerebral cortex in large numbers, mainly in deep layers and near the pial surface (**Supplementary Fig. 6e**), whereas Ascl1-derived cells were to be found numerous in the striatum but sparse in the developing neocortex (**Supplementary Fig. 6f**), with most of the cells displaying an interneuron-like morphology as expected from previous work ^53–55^. These results show that the expanding Aldh1L1+ domain generates cells with a spatial distribution distinct from the Ascl1 lineage throughout brain development despite the apparent partial overlap of Aldh1L1 and Ascl1 domains in the dLGE at E13.5. To further characterise E13.5 Aldh1L1-expressing cells, we assessed their proliferative status using co-immunostaining for cell-cycle markers, including Ki67 and PH3 (**Fig. 6f**). At E13.5, 13.73 ± 2.13 % of Aldh1L1 positive cells were actively cycling (Ki67 positive), and 11.86 ± 2.94 % undergoing mitosis (PH3 positive), indicating that only a small fraction of E13.5 Aldh1L1-expressing cells remain mitotically active at this developmental stage (**Fig. 6g**). Together, our data indicate that Aldh1L1-expressing cells at E13.5 constitute primarily a quiescent radial glia-like progenitors localised within a complex microdomain of the ventral forebrain, which includes cells expressing characteristic markers of both ventral pallium and subpallium identity.

**Fig. 6.**
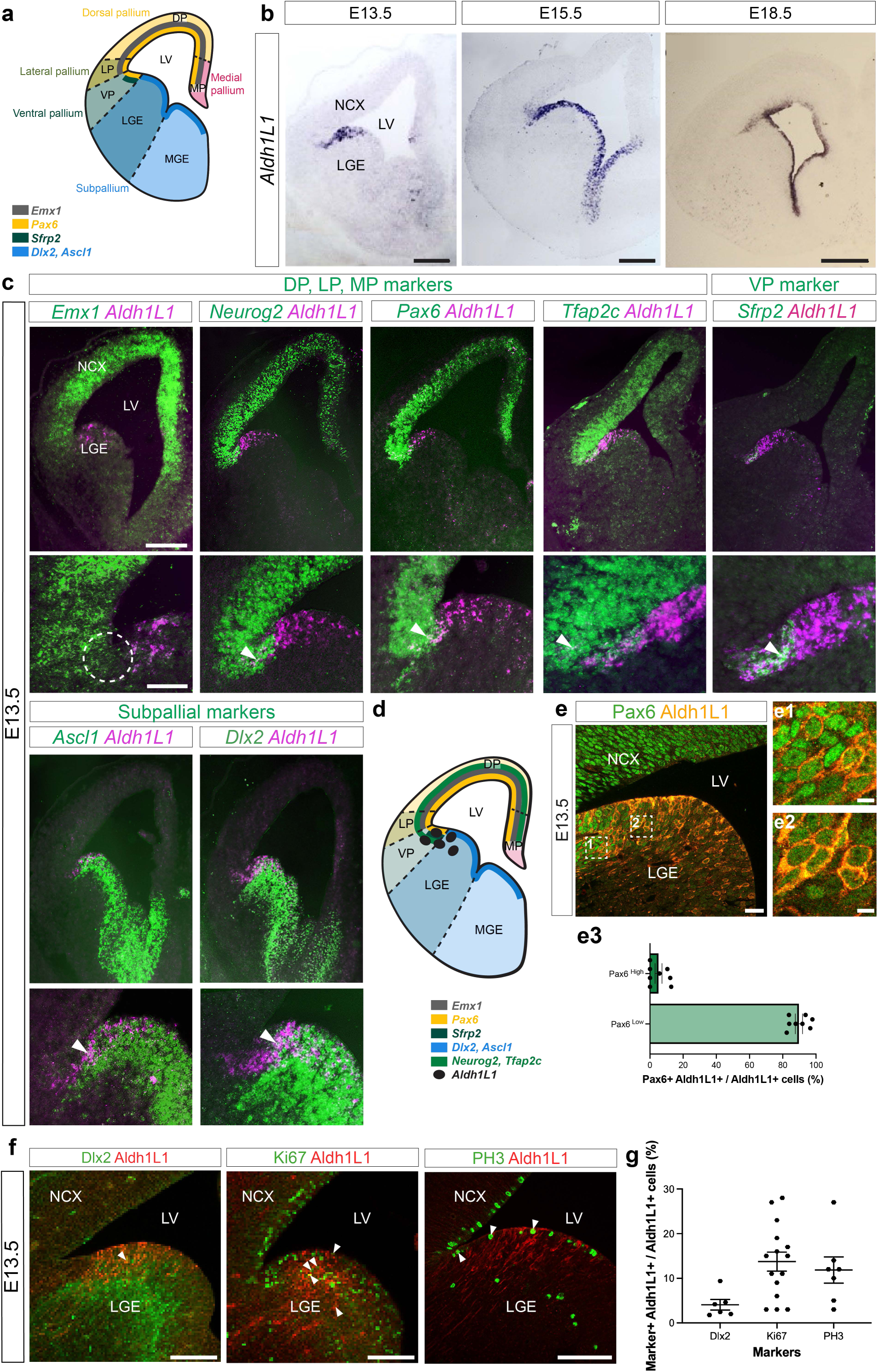
E13.5 Aldh1L1-expressing progenitors occupy a complex embryonic domain encompassing ventral pallium and subpallium but distinct from the dorsal pallium. **a** Diagram illustrating the various combinatorial gene expression domains and physical boundaries of an embryonic (E)13.5 embryonic mouse hemisphere, inspired by Modo et al 2019^69^. b *Aldh1L1* in situ hybridisation conducted at E13.5, E15.5, and E18.5 on wild-type (WT) coronal brain sections. **c** Sequential in situ hybridisation of *Aldh1L1* (purple) and recognized markers (green) for dorsal, lateral, and medial pallium (*Emx1*, *Neurog2*, *Pax6, Tfap2c*), ventral pallium (*Sfrp2*), or subpallium (*Ascl1*, *Dlx2*) on an E13.5 wild-type brain, accompanied by a summary diagram. **d** Summary diagram of the Aldh1L1 domain spanning ventral pallial and subpallial domains but located outside of the Emx1+ dorsal pallium region. **e** Confocal images depicting Aldh1L1 (orange) alongside Pax6 (green) immunostaining conducted on C57BL6 brains at E13.5. Two distinct populations of Aldh1L1-expressing cells were identified: one exhibiting high Pax6 levels (**e1**) and another exhibiting low Pax6 levels (**e2**). **e3** Quantification of Aldh1L1+ cells expressing high and low levels of Pax6 at E13.5. N = 4 independent animals, n = 8 sections. **f** Confocal images illustrating few Aldh1L1+ cells co-expressing Dlx2, Ki67 and PH3 after immunolabelling performed on coronal E13.5 C57BL6 brain sections. **g** Quantification of Aldh1L1+ progenitors co-expressing Dlx2, Ki67 and PH3 at E13.5. N = 2 (Dlx2), 4 (Ki67), and 3 (PH3) independent animals, n = 6 (Dlx2), 15 (Ki67), and 6 (PH3) sections. Dlx2 = Distal-less homeobox2, LGE = lateral ganglionic eminence, LV = lateral ventricle, MGE = medial ganglionic eminence, NCX = neocortex, PH3 = Phospho-Histone 3. Scale bar: 250 µm (**b**, **c** top); 100 µm (**c** bottom, **e**, **f**); 20 µm (**e1**, e2).

### Transcriptomic analysis of E13.5 pallial-subpallial boundary domain confirms the complex molecular signature of Aldh1L1-expressing cells, which is distinct from that of dorsal pallium progenitors, and corroborates the astroglial trajectory of their descendants

To characterise the molecular signature of Aldh1L1-expressing cells with single-cell resolution, we generated two single-cell RNA sequencing (scRNA-seq) datasets from the microdissected pallial-subpallial boundary of E13.5 C57Bl6 embryos (**Fig. 7a**). Based on established marker genes for the major developing brain cell classes **(Supplementary Table 1)**, we computed cell-type scores, annotated the integrated dataset, and identified the main cell populations, including excitatory and inhibitory neurons, radial glial cells (RGCs), and transitional populations **(Fig. 7b, Supplementary Fig. 7)**. We next subsetted RGCs regressed out cell-cycle effects, and reclustered them in SPRING space. To assess the pallial versus subpallial identity of the Aldh1L1 domain **(Fig. 7c)**, we and calculated a dorso– ventral (DV) score for each RGC based on its SPRING2 coordinate (**Fig. 7d**). RGCs were then partitioned into six DV groups using partitioning around medoids (PAM) clustering of smoothed expression profiles for significantly differentially expressed genes (**Fig. 7e**). The resulting groups were defined by the graded expression of dorsal, lateral and medial pallial markers (*Emx1*, *Pax6*, *Neurog2* and *Tfap2c*; **Fig. 7f**), ventral pallial markers (*Dbx1* and *Sfrp2*; **Fig. 7g**), and subpallial markers (*Ascl1*, *Dlx2* and *Gsx2*; **Fig. 7h**). Together, the DV score, bin assignment, and heatmap of smoothed gene expression across bins delineated six molecularly distinct progenitor domains along the DV axis **(Fig. 7g–i)**. Consistent with our ISH data (**Fig. 6c**) and despite its small size, the E13.5 Aldh1L1-expressing domain extends over bins 2 and 3. It overlaps mostly with *Sfrp2*-expressing bins, yet remains distinct from *Dbx1* (**Fig. 7g**), whose very low expression is most likely due to the late stage of embryonic development (E13.5). Furthermore, although subpallial markers such as *Ascl1*, *Dlx2* and *Gsx2* (**Fig. 7h**) were found expressed, albeit weakly, in bins 2 and 3, Pax6 is also present in these bins. The other pallial markers, such as *Neurog2*, *Tfap2c* and *Emx1* are absent (**Fig. 7f**). In conclusion, the E13.5 Aldh1L1+ domain thus displays a mixed molecular identity, encompassing transcriptomic signatures of both ventral pallial and subpallial progenitor domains, but highly distinct from the molecular identity of the dorsal pallium, confirming our results obtained with double ISH (**Fig. 6c**). To define the transcriptional program of the Aldh1l1+ population and examine its developmental progression, we integrated our E13.5 pallial–subpallial boundary dataset with a published E16.5 ganglionic eminence scRNA-seq dataset. The integrated dataset contained excitatory neurons, interneurons, intermediate progenitor cells, and RGCs; the latter were further assigned to the six DV domains defined above (**Fig. 7k**). The subsetted E13.5 and E16.5 RGCs occupied connected but partially distinct regions of the integrated embedding, and pseudotime analysis revealed a continuous developmental axis across these cells (**Fig. 7l**). Mapping RGC and astrocyte identities onto this trajectory revealed a progressive transition from progenitor-like to astrocyte-associated states (**Fig. 7m**). Consistent with this progression, gene-module analysis identified dynamic transcriptional programs across the E13.5–E16.5 continuum (**Fig. 7n**). These included progenitor and regional identity programs marked by genes such as *Gsx2, Ascl1, Id2, Pax6, Tfap2c, and Sfrp2*, as well as later glial and astrocyte-associated programs including *Fabp7, Id1, Id3, Nfia, Aldh1L1, and Aldoc*. Together, these analyses identify an Aldh1L1+ progenitor population with a molecular identity distinct from that of dorsal pallial RGCs and place these cells along a developmental continuum towards an astrocyte-associated transcriptional state. Combined with our lineage-tracing data, these findings support the conclusion that this population represents a previously unrecognised embryonic source contributing to neocortical astrocyte diversity.

**Fig. 7.**
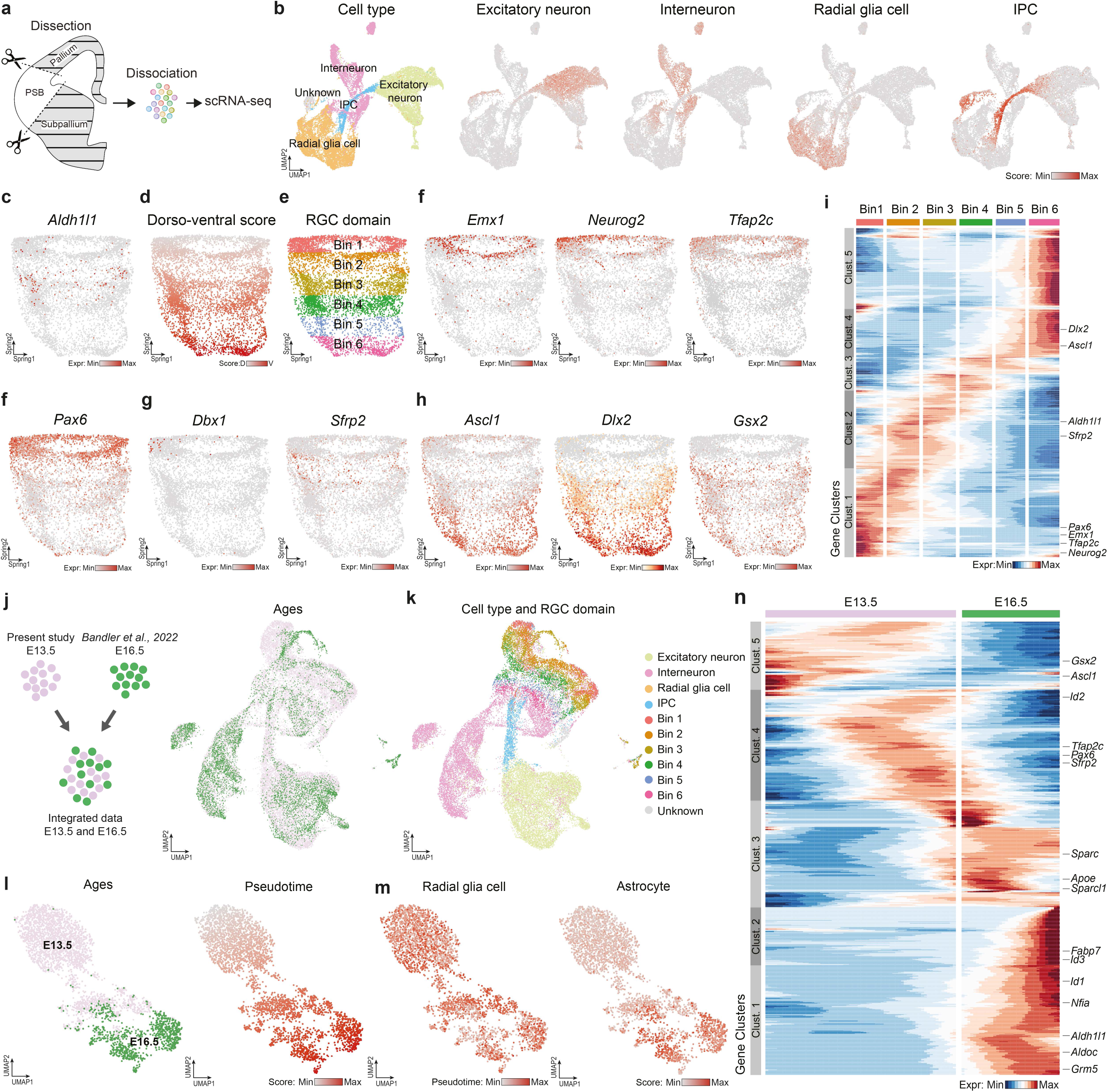
Single-cell transcriptomic profiling of the pallial-subpallial boundary (PSB) cells harvested in wild-type E13.5 embryos. **a** Schematic of pallial–subpallial boundary (PSB) dissection at embryonic day (E)13.5 and single-cell library preparation for scRNA-seq analysis. **b** UMAP representation of 23,448 single cells from the E13.5 PSB showing annotated cell types (left). Cell-type scores (right) were calculated based on established marker genes for major neocortical cell classes (full gene list in **Supplementary Table**). **c–f** SPRING representations of **subsetted radial glial cells (from panel b)** showing anatomical gene expression patterns at E13.5 after regression of cell-cycle effects. **d** SPRING representation of **subsetted radial glial cells** (from panel b) showing dorso–ventral score. **e** Identification of six radial glia cell (RGC) domains based on dorso–ventral transcriptional scoring. Shown are the expressions of the astroglial marker *Aldh1L1* (**<u>c</u>**), dorsal, lateral and medial pallium markers (*Emx1, Neurog2, Tfap2c, Pax6*) (**f**), ventral pallium markers (*Dbx1, Sfrp2*) (**<u>g</u>**), and ganglionic eminence markers (*Ascl1, Dlx2, Gsx2*) (**<u>h</u>**). **i** Heatmap showing expression of marker genes defining dorso- ventral transcriptional domains within the E13.5 PBS; with representative genes highlighted. **j** Schematic of dataset integration (left) and UMAP representation (right) of the integrated dataset, showing cells from the present study (E13.5, pink) and Bandler et al., 2022^56^ (E16.5, green). **k** UMAP visualisation showing annotated cell types and radial glia domains (RGC; Bin 1 to Bin 6). **l** UMAP plots of radial glia cells from E13.5 and E16.5, color-coded by developmental age (left) and pseudotime (right). **m** UMAP plots of radial glia cells showing radial glia cell (left) and astrocyte (right) scores calculated using established cell type marker genes (full gene list in **Supplementary Table**), illustrating progressive maturation from radial glia cells to astrocytes. **n** Heatmap of key marker genes highlighting transcriptional changes underlying the developmental transition from E13.5 to E16.5; with representative genes indicated.

## DISCUSSION

Altogether our data provide evidence of the existence of a discrete subpopulation of neocortical astrocytes derived from ventral Aldh1L1-expressing embryonic progenitors and display developmental trajectories that contrast with those of neocortical astrocytes generated from Emx1+ dorsal pallium progenitors. These two ontogenetically distinct populations of neocortical astrocytes differ in their spatial distribution across neocortical regions and in their arborisation complexity, while displaying similar territorial volume during the first postnatal weeks (**Fig. 8**).

**Fig. 8.**
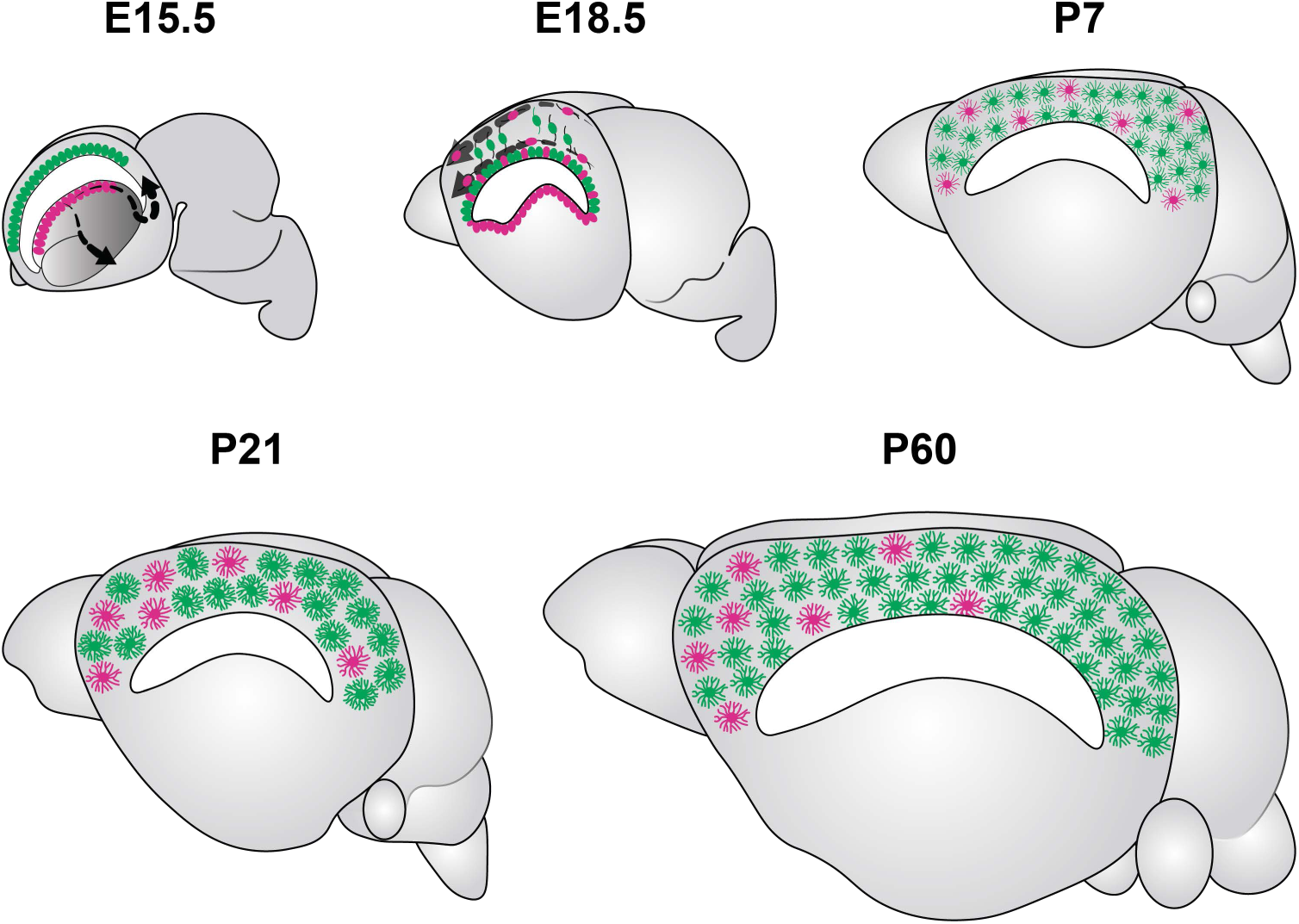
Model of divergent developmental trajectories of ontogenetically distinct neocortical astrocytes. At embryonic day (E)15.5, cells deriving from the E13.5 Aldh1L1-expressing ventral domain that line the lateral ventricle are illustrated in magenta, whereas the conventional source of cortical astrocytes, the dorsal pallium Emx1^+^ progenitors, appears in green. The progeny of E13.5 Aldh1L1-expressing progenitors migrate towards the developing neocortex via lateral and medial tangential pathways (black arrows) and along the lateral ventricles, following pial (lateral) and periventricular (medial) migratory routes (grey arrows). The descendants of E13.5 Aldh1L1^+^ embryonic progenitors progressively populate the neocortex, demonstrating a biased, region-specific distribution towards the rostral and lateral cortical areas, which becomes evident from P21 up to P60. Moreover, these ventrally derived astrocytes display a simpler morphology while maintaining, up to P21, a similar territorial volume compared to neocortical astrocytes originating from the dorsal pallium. At P60, ventrally derived astrocytes have a smaller territorial volume than dorsally derived astrocytes, yet they display comparable arbour complexity.

Even though the potential contribution of the ventral forebrain embryonic domain to the generation of cortical astrocytes has been hinted at in previous studies ^57^, additional work using straightforward genetic tools has demonstrated the regional production of astrocytes in both the spinal cord and the brain ^32^. Indeed, while the fate mapping of cells derived from the Dlx2 ventral domain revealed some GFAP+ astrocytes derived from this lineage in the neocortex, no further information was available in these studies regarding their precise final location in terms of cortical areas or layers, and most of them were actually found in the postnatal SVZ ^35^. This ventral source of neocortical astrocytes was subsequently dismissed as negligible when Tsai and collaborators used multiple transgenic Cre animals and retroviral vector injection to specifically target distinct embryonic domains and analyse the final location of their astrocyte progeny ^32^. They showed that astrocytes arising from any given embryonic domain were found only in the brain region delineated by the radial glial cells producing cells of that region, supporting a regional allocation for astrogliogenesis. Regarding the neocortex, no astrocytes were found to derive from ganglionic eminences at any developmental stages ^32^. Together with other studies confirming the production of neocortical astrocytes from radial glia cells located in the dorsal pallium that also give rise to cortical excitatory neurons ^20,21,50,58^, these data demonstrated that neocortical astrocytes originate primarily, if not exclusively, from Emx1+ cortical progenitors. Interestingly, Tsai and collaborators used the *Dbx1-Cre* transgenic mouse line ^59^ to trace the progeny of the dLGE/ventral pallium domain, and no astrocytes were found in the neocortex ^32^. The analysis of *Dbx1* expression in our scRNA-seq dataset shows that Dbx1 is not expressed in the E13.5 Aldh1L1-expressing embryonic progenitors located in the dLGE/CGE. Furthermore, no MM+ protoplasmic astrocytes were found in the neocortex of *Aldh1L1CreER^T2^;MM* animals when tamoxifen was administered before E13.5 (**Supplementary Fig. 2i**), these data explain why the production of neocortical astrocytes outside the dorsal pallium was not detected using the *Dbx1-Cre* genetic fate mapping approach. Could Aldh1L1+ embryonic progenitors correspond to the Dlx2+ subpallial cells that generate cortical astrocytes, as previously reported ^33–35^Our in situ hybridisation experiments reveal that Aldh1L1+ domain overlaps the Dlx2+ / Ascl1+ subpallial domain. However, neocortical astrocytes were rarely observed among the progeny of *Ascl1-CreER^T2^;MM* animals, at any analysed stage (**Supplementary Fig. 6**). In addition, our E13.5 scRNA-seq dataset shows that *Dlx2* and *Aldh1L1* are not co-expressed in the same cells (**Fig. 7**). Our hypothesis is that at E13.5, Aldh1L1 is mostly expressed by quiescent radial glial cells whose progeny will account for a subset of Dlx2/Ascl1 cells. However, the combination of in situ hybridisation, scRNA-seq, immunostaining and genetic fate mapping strategies we used clearly indicate that most E13.5 Aldh1L1+ cells express Pax6, albeit mostly at a low level, and Sfrp2 (**Fig. 6**), whilst not expressing Emx1, which together form a key marker combination of the embryonic ventral pallium domain ^60,61^. Interestingly, additional work supports the existence of a ventral embryonic source for neocortical astrocytes. Recent transcriptomic studies have reported the existence of clonally-related neocortical astrocytes and GABAergic interneurons in the developing cerebral cortex when viral barcoding particles were injected into the embryonic mouse at E11-E12 ^56^. Furthermore, whilst characterising two distinct lineages for neocortical astrocytes using a combination of single-cell and spatial transcriptomics, Zhou and collaborators demonstrated in *Emx1-Cre::ROSA^nT-nG^* animals at P7 the presence of approximately 30% of Sox9+/GFP-/RFP+ cells, which correspond to cortical astrocytes not derived from the Emx1 lineage ^29^. Taken together, these studies confirm the existence of an additional source of neocortical astrocytes outside the dorsal pallium, although, until our current work, they had not precisely localised these gliogenic progenitors during embryogenesis.

Our robust fate-mapping strategy enabled us to trace back the site of origin of a subset of neocortical astrocytes to the ventral pallium (VP), which was unexpected for several reasons. Firstly, despite long-lasting debate and conflicting studies about the contribution of the ventral pallium to neocortical neurons ^60^, recent studies relying on *in utero* electroporation of integrative vectors indicate that VP radial glial cells give rise to excitatory neurons that will only settle in the amygdala, olfactory cortices (insular and piriform cortex), and claustrum, and not in neocortical areas ^62^. Even though astrocytes derived from cortical progenitors located in the dorsal pallium can disseminate over several hundred micrometres and in a highly heterogeneous manner ^20^, they were known to be generated from radial glial cells that have previously produced excitatory neurons that will migrate radially toward the cortical plate and settle in a columnar manner ^32,50,58^. If these principles applied to the progeny of Aldh1L1+ radial glial cells located in the VP, astrocytes should be found exclusively in the amygdala, olfactory cortices (insular and piriform cortex) and claustrum. However, we observed that neocortical astrocytes derived from Aldh1L1+ embryonic progenitors were scattered throughout several cortical areas. This indicates that the VP does indeed generate astrocytes that settle in neocortical areas, and that it differs from the dorsal pallium in terms of the timing of astrogliogenesis. This finding is even more interesting given the peculiar evolutionary status of this embryonic structure. In theory, given that the VP is considerably more developed in sauropsids, one would expect to observe a higher proportion of VP-derived astrocytes (Emx1-lineage) in the brains of non-mammalian species compared with astrocytes derived from the dorsal pallium (Emx1+ lineage). Although care must be taken to separate molecular signatures from lineage considerations, it is therefore particularly interesting to note that the molecular trajectories of astrocytes extracted from publicly available scRNA-seq datasets, derived from reptiles ^63^ and birds ^64^ reveal that their astrocytes primarily belong to the non-Emx1-derived trajectory described by Zhou et al. ^29^. Although it is enlarged in sauropsids and considerably reduced in mammals, the VP appears to be well conserved across species, including in humans ^60^. Therefore, we would expect to find evidences of the Aldh1L1+ embryonic population and their neocortical astrocyte progeny in the human neocortex. Indeed, a population of astrocytes precursors is found in the mantle zone and subplate/intermediate zone of human foetal frontal cortical areas that are do not derive from dorsal radial glia cells but from ventral germinative zones ^65^. In addition, ALDH1L1-immunoreactive cells are present in the human developing cortex starting from the midpoint of the early foetal period onward and expand from VZ and SVZ of dLGE around 19 GW to cortical intermediate zone, subplate, and cortical plate until 29 GW ^44^. The emerging bilaminar pattern formed by one band in the marginal zone/upper cortical plate and another in the subplate/intermediate zone at the perinatal stage of early astrocytic distribution in the human medial cortex (38-39 GW; GFAP immunostaining) ^66^ resembles the lateral and medial tangential migratory pathways undertaken in mice by the VP-derived progeny we reported in our work (**Fig. 2**). Altogether, these observations support the existence of another embryonic source for neocortical astrocytes located in the ventral pallium and well-conserved across species.

By tracking the descendants of these progenitor cells throughout corticogenesis using in situ genetic fate mapping strategies, we identified key features of this vA subpopulation of astrocytes derived from the VP and compared them with neighbouring astrocytes originating from the dorsal pallium. Although distinct developmental trajectories were expected given their separated sites of origin and the longer migratory distances required to reach the developing neocortex, we found VP-derived astrocytes unevenly distributed across cortical areas. Along with a point of entry located in posterior cortical areas during late embryogenesis (**Fig. 2, Supplementary movies 1 and 2**), their preferential location in rostral cortical areas at all analysed postnatal ages (**Fig. 3**) suggests that this subpopulation of astrocytes colonises the cerebral cortex from the caudal to the rostral regions during corticogenesis. It also indicates that their dissemination across cortical areas and layers may follow a non-stochastic pattern, distinct from dorsally-derived neocortical astrocytes ^20^, while still coming into contact with blood vessels and maturing at a similar pace, as indicated by Aquaporin 4 immunolabelling (**Fig. 5**). When comparing volume and arborisation complexity between the neocortical astrocytes derived from ventral (vA) versus dorsal (dA) embryonic origins, we observed drastic differences. 3D reconstructions and detailed morphological analyses showed that vA display a simpler arborisation than dA at P21, while exhibiting a similar volume (**Fig. 4**). Identical comparative analyses were performed at P60 to determine if the simpler morphological complexity exhibited by vA could result from a delayed maturation. Our P60 data indicate that the vA and dA subpopulations no longer differed in arborisation complexity, because dA decreased their morphological complexity between P21 and P60 (**Fig. 4**). To the best of our knowledge, this age-dependent astrocyte maturation based on arborisation simplification has only been hinted at in the hippocampus ^67^ and could be associated with specific functional consequences. Together with the recently published work highlighting the critical functions of astrocyte subdomains, named leaflets at synaptic ^14^ and endfeet at vascular ^8,15^ interfaces, our data point to the importance of analysing the fine morphology of astrocytes in 3D reconstructions using semi-automated segmentation and dedicated software, rather than relying solely on 2D measurements to compare astrocyte morphologies under physiological and pathological conditions. In addition, our data indicate that astrocyte morphological maturation during cortical development is more complex than expected and continues beyond P21 with a refinement reminiscent of neuronal pruning. Although molecular cues underlying astrocyte morphogenesis during early postnatal development have been extensively studied and their multiple cellular sources reported, such as blood vessels ^8^, neurons ^18^, and astrocytes themselves relying on neuroligins ^17^ and protocadherins ^30^, the signalling pathways involved in the refinement of astrocyte arborisation after P21 have yet to be determined.

Our work uncovers the existence of a subpopulation of neocortical astrocytes that differ from those derived from dorsal pallium progenitors in morphological complexity, but not in terms of their territorial volume or their vascular contacts. Given that no obvious distinctive features have yet been identified at the level of neuronal or vascular interfaces, we can only hypothesise about a wide range of specific functions that this subpopulation of astrocytes might be particularly associated with; these will need to be carefully evaluated in upcoming studies. However, specific features of these VP-derived neocortical astrocytes may provide some clues as to their potential roles in brain functions and their potential involvement in pathologies. The first hypothesis, based on the very origin of these astrocytes, in the VP / subpallium embryonic domain, is that they would represent a remnant of ancient brain structures shared with other species, without having specific functions that differ from those of dA in the neocortex but representing an additional reservoir of astrocytes that could be mobilised in the event of a pathological condition, injury or disease. Interestingly, the Aldh1L1+ embryonic progenitors that give rise to these vA are located in the dLGE which is a structure already known to produce cortical interneurons but not excitatory neurons in the mouse brain. As these vA would share an ontogenetic relationship with interneurons, it may explain why the reprogramming of a certain fraction of neocortical astrocytes is more efficient in producing interneurons than excitatory neurons ^68^, especially if this reprogramming targets astrocytes in the most frontal regions of the neocortex, where vA can represent up to 22.12 % of the entire astrocyte population in the postnatal mouse brain. This preferential distribution of vA in frontal regions, associated with their Sparcl1/Fabp7 lineage trajectory hinted at by single-cell transcriptomic and pseudotime analyses (**Fig. 7**), suggest a potential involvement in psychiatric diseases such as schizophrenia. This would require further investigation once appropriate tools are available to selectively isolate vA from neighbouring astrocytes derived from dorsal pallium progenitors.

In conclusion, our work provides a comprehensive characterisation of a distinct subpopulation of neocortical astrocytes that is ontogenetically unrelated to the major dorsal pallium-derived astrocyte population yet coexists with them across several cortical regions in varying proportions. These astrocytes are preferentially located in frontal cortical regions and originate from Aldh1L1-expressing embryonic progenitors located in a complex zone encompassing ventral pallium and subpallium domains. Beyond their final spatial allocation and distinct origins, vA exhibit distinct developmental dynamics in arborisation and territorial volume compared with dA. Together, our findings redefine cortical astrogliogenesis as a multi-origin developmental process and establish ontogeny as a key feature of astrocyte diversity in the mammalian neocortex.

## ACKNOWLEDGEMENTS

We thank Patrick Carroll, Solène Clavreul, Jean Livet, Steven Sloan and Gerben van Hameren for scientific discussions and comments on the paper. We thank N. Tricaud and F. Ango for providing two of the transgenic mouse lines used in this study (see “Animals” section in Methods).

We acknowledge the imaging facility MRI, member of the national infrastructure France-BioImaging (https://ror.org/01y7vt929) supported by the French National Research Agency (ANR-24-INBS-0005 FBI BIOGEN) and all support provided by the technicians of the RAM-Neuro and RAM-PCEA animal facilities, and the support service of the Institute for Neuroscience of Montpellier.

## FUNDING

This work was supported by INSERM ATIP-Avenir program (PI K. Loulier), by funding from Agence Nationale de la Recherche under contract ANR-22-CE16-0012-01, by Fondation pour la Recherche Médicale (FRM ENV202109013951), by the University of Montpellier and by Fondation pour la Recherche sur le Cerveau (FRC) operation Rotary Espoir en tête 2024. RB and JZ were supported by the Swiss National Science Foundation through an Ambizione grant (PZ00P3_201995).

## AUTHOR CONTRIBUTIONS

Conceptualisation: LD, KL; Animal genotyping, histology and confocal microscope imaging, image processing and morphometric analyses: LD, DT, JD, IA, ER, PL; Data processing and analyses: LD, DT, KL; 3D multichannel multiphoton imaging: NK, HB; Transcriptomic experiments and analysis: JZ, RB; Gliovascular interface experiments and analyses: BDP, MCS; *Aldh1L1-eGFP/Olig2-dtTomato* experiments and analyses: CD, DO; *Ascl1Cre- ER^T2^;MM* lineage tracing experiments: EP; Supervision: LD, KL; Project administration: KL; Funding acquisition: KL.

## DECLARATIONS

### Ethics approval and consent to participate

All the experiments conducted on mice were approved by the local ethics committee (CEEACD/N°36) and the Ministère de la Recherche et de l’Enseignement Supérieur (authorisations #20047-2019032917332907 v2, #43098-2023031518358971 v5 and #38605-202209161547240 v5). All of the procedures were carried out in accordance with the French regulations governing animal procedures (French decree 2013–118) and with specific European Union guidelines for the protection of animal welfare (Directive 2010/63/EU).

### Competing interests

The authors declare no competing interests.

## METHODS

### Animals

All mouse procedures were approved by the local ethics committee and the French Ministry of Research and Higher Education (CEEACD/N°36, authorisations #20047-2019032917332907 v2, #43098-2023031518358971 v5 and #38605-202209161547240 v5). Mice were housed at the RAM-PCEA and RAM-Neuro animal facilities (Montpellier, France) and maintained on a standard light cycle (12 h light-dark cycle) at 21.5°C and 50% relative humidity. Water and food were provided ad libitum. MAGIC Marker mice were produced by intercrossing two transgenic mouse lines generated by pronuclear injection of the *CAG-Cytbow* and *CAG-Nucbow* transgenes described in Loulier et al, Neuron, 2014. The resulting double transgenic mice were then crossed with B6N.FVB-Tg (Aldh1L1-CreER^T2^)1Khakh/J mice ^39^ (JAX strain 031008, a gift of N. Tricaud), and their *Aldh1L1-CreER^T2^;CAG-Cytbow;CAG-Nucbow* (aka *Aldh1L1-CreER^T2^;MM*) offspring analysed at distinct developmental stages following tamoxifen (T5648; Sigma) oral administration at defined embryonic stages or subcutaneous injection at postnatal day (P)4.

Additional reporter lines were also used in this study: *Aldh1L1-CreER^T2^;Ai14* (JAX strain 007914 (B6.Cg-Gt(ROSA)26Sortm14(CAG-tdTomato)Hze/J, a gift of F. Ango), *Aldh1L1-eGFP* (JAX strain 030247 (B6;FVB-Tg(Aldh1L1-EGFP/Rpl10a)JD130Htz/J)); *Olig2-tdTomato* (MGI ID: 5311714 (Tg(Olig2-tdTomato)TH39Gsat)). Mice were mated overnight, and the day of the vaginal plug observation was defined as embryonic day (E)0.5.

### In vivo experiments

#### Tamoxifen induction

At E13.5, timed-pregnant mice received 2 mg of tamoxifen by gavage using a feeding tube (FSTx-10). At the postnatal day (P)4, pups were injected subcutaneously with 0.2 mg of tamoxifen (Sigma T5648) using a 30G needle. The tamoxifen was dissolved in corn oil.

#### In utero electroporation

At E15.5, timed-mated C57Bl/6J pregnant females were anesthetised with an intraperitoneal injection of ketamine (62.5 mg/kg) and xylazine (12.5 mg/kg), followed by a subcutaneous analgesia (buprenorphine, 0.048 µg/g). After the exposure of the uterine horns via a laparotomy, 1 µl of plasmid vector DNA mix (*PB-CAG::Cytbow* 0.8 ug/ul, Addgene ID # 158995; *Tol2-CAG::Nucbow* 0.8 ug/ul, Addgene ID # 158992; *CAG-SeCre* 0.16 ug/ul; *CAG-PBase* 0.4 ug/ul; *CAG-Tol2ase* 0.4 ug/ul; ^40^) was micro-injected into the lateral ventricle and then unilaterally electroporated into pallial progenitors lining up the dorsal wall of the ventricle using 3-mm diameter tweezer-type electrodes (3 pulses of 50 ms with an amplitude of 30 V). Upon completion of the surgery, the uterus was repositioned into the abdomen with physiological serum, and the wound was sutured with absorbable 4/0 sutures, thereby permitting normal embryonic development. The full procedure is described in detail in previous publications ^24,48^.

#### Adoption

Due to delivery complications in tamoxifen-injected pregnant mice, time-mated Swiss mice were utilised to adopt the recombined pups. Foster Swiss females were mated several days prior to the breeding of the *Aldh1L1-CreER^T2^;MM* mice. To obtain postnatal stages, tamoxifen-induced pregnant females were euthanised by cervical dislocation between E18.5 and E19.5. Embryos were obtained via caesarean section, rinsed in warm 1X phosphate-buffered saline (PBS), gently dried, and temporarily placed in a warm artificial nest containing 2-3 foster pups along with used nesting material before being transferred to the adoptive Swiss mouse.

#### Tissue collection

##### Sample collection for in situ hybridisation, immunostaining, and confocal imaging

Pregnant mice were euthanized by cervical dislocation at the designated developmental stages to harvest embryonic brains on ice. Dissected embryonic brains were subsequently fixed in a paraformaldehyde (PFA)-based fixative (Antigenfix, Diapath) for three hours at 4°C with agitation for immunostaining or in fresh 4% PFA-PBS overnight at 4°C for in situ hybridisation, prior to being washed in 1X PBS. Pups and adult mice were euthanized using EUTHASOL® VET and perfused intracardially with Antigenfix for postnatal analysis. Postnatal dissected brains were fixed in Antigenfix overnight at 4°C with agitation, followed by three 10-min washes in 1X PBS.

#### Sample collection for single cell RNA sequencing

Brains from E13.5 embryos were dissected on ice in Leibowitz medium supplemented with 5% FBS. For each embryo, a region spanning the pallial-subpallial border was microdissected from both hemispheres. Tissues were enzymatically dissociated using the Papain Dissociation System (Worthington, #LK003150) according to the manufacturer’s protocol and mechanically processed using the gentleMACS Dissociator according to the manufacturer’s instructions.

### Histology

#### Clearing of whole embryonic hemisphere

Whole embryonic hemispheres at E15.5 and E17.5 were incubated in modified CUBIC R1 reagent (34 wt% N,N,N’,N’-tetrakis(2-hydroxypropyl) ethylenediamine (Sigma 122262), 15 wt% Triton-X100 (Sigma), 51 wt% H_2_O) at 37°C under agitation until cleared.

#### Immunostaining on floating sections

Immunostaining was performed on 100 µm brain sections collected using vibrating microtomes (VT1000S, Leica; HM650V, Thermo Scientific^TM^) after embedding in 3% agarose.

Floating sections were incubated from 30 min to 1 h in blocking and permeabilization solution with 0.2% Gelatin and 0.5% Triton X-100 (Sigma) in 1X PBS at room temperature (RT), then incubated overnight with at least one of the following primary antibodies: anti-Aldh1L1 (1:800, Abcam ab87117); anti AQP4 (1:500, Merck A5971 or Alomone Labs AQP-004); anti-Dlx2 (1:100, Santa Cruz Biotechnology sc-393879); anti-CD31 (1:300, Bio-Techne AF3628); anti-GFP (1:500, Abcam AB13970); anti-Histone H3 (1:500, Abcam ab10543); anti-Ki67 (1:200, Abcam ab16667); anti-NeuN (1:1000, Millipore ABN91); anti-Olig2 (1:400, R&D Systems AF2418); anti-Pax6 (1:500, Sigma AB2237); anti-SMA-Cy3 (1:300, Merck C6198); anti-Sox9 (1:500, R&D Systems AF3075).

For the anti-Aldh1L1 immunostaining, sections were unmasked in citrate buffer (10 mM Trisodium citrate, 0.05 % Tween-20 at pH 6.0) by incubating them for 5 min at RT, 15 min at 68°C and then cooling down for 10 min at RT inside a humidity chamber. Sections were then washed, blocked and permeabilized as previously described.

After washing three times in 1X PBS, sections were incubated for 3 h at RT with specific secondary antibodies: anti-chicken Alexa Fluor 488 (1:500, Thermo Fisher Scientific A32931TR); anti-goat conjugated with Alexa Fluor 488 (1:400, Jackson ImmunoResearch 705-545-147) or 647 (1:400, Jackson ImmunoResearch 705-605-147 or 1:2500, Thermo Fisher A21447); anti-mouse Alexa Fluor 488 (1:400, Jackson ImmunoResearch 115-545- 146); anti-rabbit conjugated with Alexa Fluor 488 (1:400, Jackson ImmunoResearch 111-545-144), 594 (1:400, Jackson ImmunoResearch 111-585-144) or 647 (1: 1000, Thermo Fisher A-21245 or 1:2500, Thermo Fisher A21245); anti-rat Alexa Fluor 488 (1:400, Jackson ImmunoResearch 112-545-143). Sections were then rinsed three times with 1X PBS and mounted on slides with Vectashield Plus (Vector Labs) or fresh RapiClear® 1.49 mounting media.

#### Cryosections for in situ hybridisation and double constitutive reporter approach

Washed embryonic brains post-fixed overnight were then cryoprotected in 25% sucrose overnight at 4°C. They were embedded in OCT Compound (Tissue-Tek) in inclusion molds and stored at -70 to 80°C. Coronal cryosections 14/15 µm thick were obtained with a cryostat (Microm/ Leica CM9150), mounted on SuperFrost + slides (VWR) and stored at -20°C.

##### Preparation of probes labelled with Digoxigenin or Fluorescein

Linearised plasmids containing the cDNA of *Aldh1L1* (*Source Bioscience*), *Ascl1* (F. Guillemot), *Dlx2* (E. Bellefroid), *Neurog2* (F. Guillemot) and *Pax6* (J. Ericson), and PCR products for *Emx1* (Forward: 5’-TGGTGGCCAAGGATGGTGGC-3’ / Reverse: 5’-AATTAACCCTCACTAAAGGGAGGCGGTGGCCAAAGAAGCGAT-3’), Sfrp2 (Forward: 5’-CATTTACAAGCTGAACGGCG-3’ / Reverse : 5’-AATTAACCCTCACTAAAGGGAGTGTACAAGATCTGGATGGGC-3’) and Tfap2c (Forward : 5’-AGGTGTGCAAGGAGTTCACT-3’ / Reverse : 5’-AATTAACCCTCACTAAAGGGAGGAACCAAACCAAGTGGCTTC-3’) were used as templates for the synthesis of antisense RNA probes with T3, T7 or SP6 RNA polymerases, using the Digoxigenin or Fluorescein Labelling Kit, as recommended by the supplier (Roche). Probes were purified on G50 columns (GE Healthcare) and tested on agarose gel.

##### Solutions used for in situ hybridisation

5X MAB: 58 g maleic acid; 44 g NaCl; H_2_O qsp 800 mL; pH 7.5 with NaOH pellets; qsp 1 L; autoclave.

10% blocking solution: 10 g of blocking reagent (Boehringer 1096176) in 100 mL maleic acid solution.

10X salt: 11.4 g NaCl; 8 mL 1 M Tris-HCl; 2 mL 1M Tris-base; 780 mg NaH_2_PO_4_, 2 H_2_O; 710 mg NaHPO_4_; 10 mL 0.5 M EDTA; H_2_O qsp 100 mL.

Buffer B3: 24.2 g Tris pH 9.5 (100 mM); 11.68 g NaCl (100 mM), 2 L, pH 9.5; autoclave; add 0.51 g MgCl_2_/500 mL; 0.1% Tween-20; 2 mM levamisole.

Hybridisation buffer: 1 mL 10X salt; 5 mL formamide; 2 mL 50% dextran sulphate; 1 mL yeast tRNA 10 mg/mL; 200 µL Denhardt’s 50X; 800 µL H_2_O DEPC.

Maleic acid solution: 1.16 g maleic acid (Fluka 63180); 877 mg NaCl; make up to 100 mL; pH 7; dissolve solution 2 to 3 min in microwave; autoclave; make 1 mL aliquots and store at - 20°C.

##### Sequential double in situ hybridisation

Sections were thawed for 30 min at RT. Digoxigenin- and Fluorescein-labelled antisense RNA probes were diluted together (each 1:150) in hybridisation buffer, denatured 10 min at 70°C and then cooled on ice. 300 µL of this probe mixture was applied on each slide under coverslips, then incubated overnight at 70°C in a humidified chamber with a solution of “50% Formamide, H_2_O”. Slides were rinsed for 10 min in wash buffer (50% Formamide, 1X SSC, 0.1% Tween-20) in a 70°C tray to remove coverslips and excess probes, followed by two 45-min washes in the same buffer at 70°C. After two 10 min rinses in “1X Maleic Acid pH 7.5, 0.1% Tween-20” (MABT) buffer at RT, slides were blocked for 1 h in “MABT, 20% sheep serum, 2% blocking reagent” (Roche) at RT. The anti-Digoxigenin antibody coupled to Alkaline Phosphatase (1:2000 in blocking buffer, Roche 1093274) was incubated for 2 h at RT. After three washes of 10 min in MABT at RT, the slides were rinsed 3 x 10 min in 0.1M Tris pH 8.2 at RT and incubated between 12 h and 36 h in “Fast Red” (Sigma), an Alkaline Phosphatase substrate, previously prepared according to the supplier’s recommendations and filtered through a 0.22 µm filter. After satisfactory staining (visible red staining and fluorescence), the sections were washed for 30 min in 0.1M Tris pH 8.2 and temporarily mounted under coverslips in “90% Glycerol, 0.1 M Tris pH 8.2” for imaging. Red labelling images were acquired in transmitted light and/or fluorescence using a Zeiss Microscope. The slides were then rinsed 10 min in 0.1 M Tris pH 8.2 to remove the coverslips and incubated 30 min at RT in “100 mM Glycine pH 2.2, 0.1% Tween-20” to inactivate Alkaline Phosphatase activity. After two washes in “1X PBS, 0.1% Tween-20” (PBT) of 10 min, the slides were refixed in “4% PFA-PBT” for 10 min before being washed again three times for 10 min in PBT and twice 10 min in MABT. Slides were reblocked for 1 h in “MABT, 20% sheep serum, 2% blocking reagent” (Roche) at RT and incubated with 400 µL of anti-Fluorescein antibody coupled to Alkaline Phosphatase (1:1000, Roche) under coverslips overnight at 4°C to detect the second probe. Slides were washed three times 10 min in MABT at RT and then rinsed twice 15 min in buffer B3. They were then incubated with a second Alkaline Phosphatase substrate, “NBT-BCIP” (Roche) prepared according to the supplier’s instructions in buffer B3. After satisfactory dark-blue staining (12-36 h), revelation was stopped in water. The first red stain was then dissolved by successive baths in 20%, 50% and 75% ethanol solutions for 1 min, then twice 10 min in 100% ethanol, until the red stain completely disappeared. Sections were then rehydrated into a series of baths containing 75%, 50% and 20% ethanol. Slides were then washed with water, air-dried and mounted under coverslips in aqueous mounting medium (Mowiol). Images of the blue revelation were acquired using the same Zeiss microscope at the same magnifications. Images of the two successive stains (red and blue) taken in the same region were transformed into pseudo-fluorescence and superimposed using Photoshop software.

##### Simple in situ hybridisation

The more sensitive Digoxigenin-labelled probes were diluted in hybridisation buffer and deposited on the slides overnight at 70°C. After washing in “50% Formamide, 1X SSC, 0.1% Tween-20” at 70°C, rinsing in MABT at RT and blocking in “MABT, 20% sheep serum, 2% blocking reagent” (Roche), the slides were incubated overnight at 4°C with anti-Digoxigenin antibody coupled to Alkaline Phosphatase diluted 1:2000. After washing in MABT, the slides were rinsed in buffer B3 and incubated with the “NBT-BCIP” substrate (Roche). After staining, slides were washed in water and mounted under coverslips in Mowiol. Images were acquired with a Zeiss microscope and processed with Photoshop software.

### scRNA-seq

#### scRNA-seq library construction and preprocessing

Single-cell libraries were generated using the 10x Genomics Chromium Single Cell 3′ platform with the following kits: Chromium Single Cell 3′ Library & Gel Bead Kit v3.1 (PN-1000121), Chromium Single Cell 3′ Chip Kit v3.1 (PN-1000127), and Dual Index Kit TT Set A (PN-1000215), following the manufacturer’s guidelines. Libraries were sequenced on an Illumina NovaSeq 6000 at the iGE3 Genomics Platform (University of Geneva). FASTQ files were aligned to the mm10-2.1.0 reference transcriptome and UMI-count matrices were generated using Cell Ranger (v6.0.1; 10x Genomics).

#### scRNA-seq data analyses

All scRNA-seq analyses were performed in R (version 4.2.1) using the Seurat workflow (version 4.3.0.1)1 for cell filtering and data normalization. Each dataset was imported into R as a raw count matrix and converted to a Seurat object using the CreateSeuratObject() function with the following parameters: min.cells = 3 and min.features = 200. Cells with >10% mitochondrial gene content were excluded. For clustering, we constructed a shared nearest-neighbor graph using FindNeighbors() on the Harmony-corrected embeddings (30 dimensions). Clusters were identified with the SLM algorithm via FindClusters() (dims = 30, resolution = 0.5). Cell-type scores were computed using known brain cell-type markers via the CellCycleScoring() function. Cluster identities were assigned manually based on marker expression, cell-type scores, and reference datasets from the mouse brain atlas (http://mousebrain.org)3. Clusters that could not be confidently annotated were labelled as “Undefined”.

For plotting gene expression at the single-cell level, raw gene counts were normalized by total counts per cell, multiplied by 10,000 (to account for sequencing depth), and log-transformed using the natural logarithm (log1p) to ensure comparability across cells and to reduce technical variability.

RGC domain identification was performed according to the pipeline established in the previous study ^56^. Briefly, RGCs were embedded using SPRING dimensionality reduction, which accurately captured the DV axis. Dorsal identity was defined by markers Pax6 and Neurog2, whereas ventral identity was defined by Gsx2 and Ascl1. The DV score of each cell was derived from its SPRING2 coordinate. RGCs were then partitioned into six equally sized DV groups using partition-around-medoids (PAM) clustering on the smoothed expression profiles of significantly differentially expressed genes. Genes varying along the DV axis were identified using the differentialGeneTest() function from Monocle v2.145 and grouped by hierarchical clustering.

Integration of E13.5 and E16.56 datasets was performed using Harmony (version 0.1.1) via the RunHarmony() function with default parameters. Cell-type and RGC-domain annotations were transferred from E13.5 to E16.5 using FindTransferAnchors() and TransferData().

For pseudotime analysis, RGCs were extracted from the integrated dataset and visualised in UMAP space. A principal curve was fitted onto the UMAP embedding using principal_curve() from the princurve package, and pseudotime values were assigned based on position along the curve. Genes significantly associated with pseudotime were identified using the differentialGeneTest() function from Monocle5 (v2.14) and grouped into co-expression modules by hierarchical clustering.

### Imaging

#### Confocal microscopy

Confocal images were acquired with a 20 × 0.75/0.8 NA or a 40 × 1.3 NA objective on a Zeiss LSM 880 or on a Stellaris 5 confocal microscope. To image MAGIC Markers, endogenous mCerulean/mTurquoise2, EYFP and tdTomato/mCherry fluorescence was sequentially excited using 440, 515 and 559 nm laser lines, or a white-light laser supplemented with an additional 440 nm laser line. Each fluorophore used for the immunostaining (Alexa Fluor 488, 594, and 647) was excited by the 488, 561, and 633 nm laser lines, respectively. XY resolution and Z step size were automatically optimized based on the Nyquist criterion. Images were acquired with a resolution of 1024 × 1024 pixels.

#### Multiphoton microscopy for whole embryonic hemisphere imaging

A custom imaging chamber was assembled by stacking PTFE layers (Sigma Z278769-1EA) on a glass slide to adjust the chamber size to each cleared hemisphere. The chamber was filled with modified R1 clearing reagent to maintain optical transparency. The medial face of the cleared hemisphere was glued on the 0.17-mm thick coverslip and immersed into the chamber solution. An additional PTFE layer was added on the top of the coverslip top to ensure stable water immersion of the objective and thus long-duration imaging.

For simultaneous and volumetric imaging of three fluorescent proteins expressed by MAGIC Markers mouse, we used a LSM 7 MP OPO multiphoton microscope (Zeiss, France) coupled to an Axio Examiner Z.1 upright microscope. The imaging system was equipped with a tunable Ti:Sapphire (T:S) femtosecond laser, operating from 680 to 1080 nm (140 fs pulses, 80 MHz; Chameleon Ultra II, Coherent, France). This laser pumped from an optical parametric oscillator (Chameleon Compact OPO, Coherent, France), providing additional excitation wavelengths ranging from 1050 to 1300 nm (200 fs pulses, 80 MHz). Timed synchronisation of these two pulsed lasers was used for tricolour excitation. For optimal two-photon excitation of mTurquoise2/mCerulean and tdTomato/mCherry, two separate wavelengths at λ_T:S_ =850 nm and λ_OPO_ =1100 nm were used, respectively. To generate a “virtual” wavelength for two-photon excitation of EYFP (λ_3_ = 959 nm), we used the frequency mixing technique achieved by time-synchronizing the excitation beams λ_T:S_ = 850 nm and λ_OPO_ = 1100 nm (Δt = 0) with λ = 2/(1/λ_T:S_+1/λ_OPO_) using a delay line. The two beams were also spatially synchronised using a periscope in the XY plane and a collimator in the Z direction ^38^. Band-pass filters were used in front of the detectors to simultaneously collect blue (475/64 nm), green (542/50 nm) and red (570-610 nm) fluorescence. The refractive index of the objective was adjusted to the imaging medium. Using a XY motorized stage, we performed complete imaging of each hemisphere covering about 15 x 15 fields, depending on the sample size. A 20x W Plan Apochromat DIC VIS-IR objective was used with a zoom of 1.2, a z-step size of 10 µm, a pixel size of 0.692 µm in XY, and a pixel dwell time of1.58 µs. The “Stitching” module of the ZEN software was used to assemble the mosaic images using the green channel as reference.

### Analysis

#### 3D analysis

3D image visualization was performed on Imaris software (Oxford Instruments).

Astrocyte morphology from frontal and somatosensory cortical regions was assessed by combining volumetric and arborisation analyses using Imaris and Vaa3D (version x.1.1.2), respectively, as previously described ^24^.

#### Cell counting

Cell counting was performed using the “Cell Counter” plugin in ImageJ (version 1.54f). Quantification was performed over 600 µm x 600 µm area in 3 distinct cortical regions (frontal, medial, caudal) and subdivided into upper and lower layers based on the cortical thickness midpoint or the entire cerebral cortex.

#### Statistics

Statistical analysis and graphs were carried out using GraphPad Prism. The normality of the data distribution was checked using the Shapiro-Wilk statistical test. When the data were normally distributed (p > 0.05), parametric tests were used (multiple t-tests, 2-factor ANOVA). When the data distribution did not follow a normal distribution (p < 0.05), non-parametric tests were used (Mann-Whitney or Kruskal-Wallis). A Welch correction was applied when variances differed significantly between groups (Brown-Forsythe test, p < 0.05). Data were represented in a scatter plot with the standard error of the mean.

## SUPPLEMENTARY MATERIALS

### SUPPLEMENTARY FIGURE LEGENDS

**Supplementary Fig. 1**. *Postnatal and embryonic Aldh1L1 lineage tracing strategies revealed distinct contributions to the brain cells and regions at P21.* **a** Representative sagittal section from a P21 *Aldh1L1-CreER^T2^;MM* mouse (TMX at P4) showing widespread postnatal labelling throughout several forebrain regions. **b** Sagittal section from a P21 *Aldh1L1-CreER^T2^;MM* mouse (TMX at 13.5) demonstrating that the E13.5 Aldh1L1-expressing progenitor domain gives rise to distinct regions, including ventral pallial derivatives (piriform cortex, amygdala and claustrum, highlighted in white) and subpallial derivatives (caudate putamen and pallidum, indicated in yellow). **c, d, e, f, g**. Cell fate of Aldh1L1 progenitors tagged at E13.5 and analysed at postnatal days (P)21/22. This representative image demonstrates the diversity of cell types generated from E13.5 Aldh1L1 progenitors including various glial and neuronal cell types within the postnatal forebrain: cortical oligodendrocytes (**c**), cortical interneurons (**d**), confirmed via immunostaining for GAD67 (**d’**), pial astrocytes (**e**), protoplasmic astrocytes (PrA) within the cortical parenchyma (**f**), and fibrous astrocytes located in the corpus callosum (**g**). CLA= claustrum, PrA = protoplasmic astrocyte. Scale bars: 500 µm (a, b); 20 µm (c, d, d’, e, f, g).

**Supplementary Fig. 2**. *Developmental timing of Aldh1L1 expression during embryogenesis labelling influences the proportion of protoplasmic astrocytes generated in the postnatal neocortex.* **a-d** Representative confocal images of the Aldh1L1 progenitor domain at the pallium-subpallium boundary at the time of tamoxifen administration at E12.5 (**a**), E13.5 (**b**), E14.5 (**c**), and E15.5 (**d**), obtained via immunostaining revealing Aldh1L1 expression. **e-h** Representative confocal images of the neocortex of *Aldh1L1-CreER^T2^;MM* mice at P21/22 following tamoxifen-induced recombination at E12.5 (**e**), E13.5 (**f**), E15.5 (**g**) or after triple induction at E13.5, E14.5 and E15.5 (**h**). **e’-h’** Close-up of the upper cortical layers showing labelled protoplasmic astrocytes (white arrowheads) and interneurons (empty arrowheads) after various developmental stages of tamoxifen administration. **i** Quantification of protoplasmic astrocytes (PrA) among MM+ cells in the frontal and somatosensory neocortex at P21/22 following recombination at E12.5, E13.5, E15.5, or triple induction at E13.5, E14.5, and E15.5. E12.5: N = 3 animals, n = 48 regions from 8 sections; E13.5: N = 15 animals, n = 152 regions from 36 sections; E15.5: N = 2 animals, n = 23 regions from 4 sections; E13.5+E14.5+E15.5: N = 2 animals, n = 30 regions from 5 sections. Data were analysed using a Kruskal–Wallis test with multiple comparisons. E12.5 versus E13.5, p < 0.0001; E12.5 versus E15.5, p < 0.0001; E12.5 versus E13.5+E14.5+E15.5, p < 0.0001; E13.5 versus E15.5, p < 0.0001; E13.5 versus E13.5+E14.5+E15.5, p < 0.0001; E15.5 versus E13.5+E14.5+E15.5, p = 0.4844. CP = cortical plate. Scale bars: 50 µm (a-d), 500 µm (e-h), 20 µm (e’-h’).

**Supplementary Fig. 3**. *Validation of the specificity of the Aldh1L1-CreER^T2^ lineage-tracing strategy employed in this study.* **a1** Coronal section of *Aldh1L1-CreER^T2^;MM* forebrain at E15.5 without tamoxifen induction, highlighting endogenous fluorescent proteins and GFP immunostaining at the boundary between the pallium (dorsal forebrain) and subpallium (ventral forebrain). **a2** Higher-magnification view of the boxed region in a1. **a3** GFP immunostaining reveals default EBFP nuclear labelling in non-recombined MAGIC Markers reporter cells, with no evidence of Cre-dependent recombination, confirming the absence of reporter leakiness. **b** Independent validation employing a complementary transgenic approach using the *Aldh1L1-CreER^T2^;Ai14* reporter line following tamoxifen administration at E13.5. Nuclei were counterstained with Hoechst. **b1** Coronal forebrain section at E15.5 displaying tdTomato-expressing Aldh1L1-lineage cells. **b2** Forebrain coronal section at E18.5 illustrating the distribution of labelled progeny. IF = immunofluorescence, GFP = Green Fluorescent Protein. Scale bars: 200 µm (a1, b1, b2), 50 µm (a2).

**Supplementary Fig. 4.** *Developmental trajectories of the morphogenesis of astrocytes derived from the dorsal pallium (dA) compared to the astrocytes derived from E13.5 Aldh1L1-expressing ventral progenitors (vA)*. **a** Territorial volume evolution of dA vs vA at P7, P21, and P60 following TMX administration at E13.5. Two-way ANOVA with a multiple comparisons test, N = 8 (P7), 12 (P21), and 9 (P60) independent animals, n = 22 (P7), 23 (P21), and 25 (P60) cells. **b-d** Progression of the arborisation features (evolution of the number of nodes (**b**), branches (**c**) and tips (**d**)) of dA and vA astrocytes at P7, P21, and P60 following E13.5 recombination. Two-way ANOVA with a multiple comparisons test, N = 10 (P7), 11 (P21), and 7 (P60) independent animals, n = 32 (P7), 30 (P21), and 22 (P60) cells. **e** dA territorial volume between upper and deep layers at P21. Mann-Whitney test, p = 0.0173; N = 4 independent animals, n = 11 dA and 12 vA. **f** vA territorial volume between upper and deep layer at P60. Mann-Whitney test, p = 0.0238; N = 5 independent animals, n = 16 dA and 9 vA. Error bars represent means ± SEM.

**Supplementary Fig. 5.** *Aldh1L1-eGFP;Olig2-tdTomato reporter mice delineate distinct embryonic forebrain populations.* **a** Coronal section from an *Aldh1L1-eGFP; Olig2-tdTomato* embryo at E12.5 with no eGFP expressed in progenitors lining the lateral ventricle, and tdTomato solely expressed in the medial ganglionic eminence. **b** Aldh1L1-eGFP-positive cells detected in ventral forebrain regions and along trajectories extending progressively towards the developing dorsal pallium at E16.5. **c** Broad distribution of Aldh1L1-eGFP-positive cells within the developing dorsal pallium at E17.5. This double-constitutive, non-inducible reporter approach identifies cells actively expressing Aldh1L1 and/or Olig2 at the time of analysis, and confirms the results obtained with our *Aldh1L1Cre-ER^T2^;MM* lineage strategy. CP = cortical plate, LV = lateral ventricle. Scale bars: 100 µm (**a**, **b**, **c**).

**Supplementary Fig. 6.** *Distinct developmental trajectories of Aldh1L1 and Ascl1 lineages tagged at E13.5 throughout brain development.* MAGIC Marker expression driven by the Aldh1L1 promoter (**a**, **c**, **e**) or the Ascl1 promoter using *Ascl1-CreER^T2^;MM* mice (**b**, **d**, **f**) on coronal brain sections at E15.5 (**a**, **b**) and E18.5 (**c**, **d**), and on sagittal sections at P7 (**e**) and P8 (**f**), following tamoxifen induction at E13.5. CP = cortical plate, GE = ganglionic eminence, LV = lateral ventricle, NCX = cerebral neocortex, ST = striatum. Scale bars: 200 µm.

## SUPPLEMENTARY MOVIE LEGENDS

**Supplementary Movie 1.** *Multiphoton imaging of a E15.5 Aldh1L1-CreER^T2^; MAGIC Markers embryonic brain following tamoxifen injection at E13.5.* Sequential optical sections of a cleared cerebral hemisphere from *Aldh1L1-CreER^T2^; MAGIC Markers* embryo at E15.5, which received tamoxifen injection at E13.5, were acquired by multichannel multiphoton microscopy. Captured images were compiled into an animated sequence showing the three-dimensional ventral distribution of multicoloured cells derived from Aldh1L1-expressing embryonic progenitors, identified by their expression of the fluorescent proteins of the MAGIC Markers at E15.5.

**Supplementary Movie 2.** *Multiphoton imaging of a E17.5 Aldh1L1-CreER^T2^;MAGIC Markers embryonic brain following tamoxifen injection at E13.5.* Sequential optical sections of a cleared cerebral hemisphere from *Aldh1L1-CreER^T2^; MAGIC Markers* embryo at E15.5, which received tamoxifen injection at E13.5, were acquired by multichannel multiphoton microscopy. Captured images were compiled into an animated sequence showing the three- dimensional distribution of multicoloured cells derived from Aldh1L1-expressing embryonic progenitors, identified by their expression of the fluorescent proteins of the MAGIC Markers at E17.5.

## SUPPLEMENTARY TABLE LEGEND

**Supplementary Table.** *Marker genes used for cell-type annotation.* Cell types were classified according to the expression of several established canonical marker genes.

## Notes

### Competing Interest Statement

The authors have declared no competing interest.

## REFERENCES

1. Allen, N. J. Astrocyte Regulation of Synaptic Behavior. Annu. Rev. Cell Dev. Biol. 30, 439–463 (2014).

2. Allen, N. J. & Eroglu, C. Cell Biology of Astrocyte-Synapse Interactions. Neuron 96, 697– 708 (2017).

3. Khakh, B. S. & Deneen, B. The Emerging Nature of Astrocyte Diversity. Annu. Rev. Neurosci. 42, 187–207 (2019).

4. Cooper, M. L. et al. Astrocytes connect specific brain regions through plastic networks. Nature 655, 183–191 (2026).

5. Ribot, J. et al. Astrocytes close the mouse critical period for visual plasticity. Science 373, 77–81 (2021).

6. Wahis, J. et al. Astrocytes mediate the effect of oxytocin in the central amygdala on neuronal activity and affective states in rodents. Nat. Neurosci. 24, 529–541 (2021).

7. Williamson, M. R. et al. Learning-associated astrocyte ensembles regulate memory recall. Nature 637, 478–486 (2025).

8. Freitas-Andrade, M. et al. Astroglial Hmgb1 regulates postnatal astrocyte morphogenesis and cerebrovascular maturation. Nat. Commun. 14, 4965 (2023).

9. Díaz-Castro, B., Robel, S. & Mishra, A. Astrocyte Endfeet in Brain Function and Pathology: Open Questions. Annu. Rev. Neurosci. 46, 101–121 (2023).

10. Cohen Salmon, M., et al. Astrocytes in the regulation of cerebrovascular functions. Glia 10.1002/glia.23924 (2020) doi:10.1002/glia.23924.

11. Boulay, A.-C. et al. Immune Quiescence of the Brain Is Set by Astroglial Connexin 43. J. Neurosci. 35, 4427–4439 (2015).

12. Boulay, A.-C., Cisternino, S. & Cohen-Salmon, M. Immunoregulation at the gliovascular unit in the healthy brain: A focus on Connexin 43. Brain. Behav. Immun. 56, 1–9 (2016).

13. Iadecola, C. The Neurovascular Unit Coming of Age: A Journey through Neurovascular Coupling in Health and Disease. Neuron 96, 17–42 (2017).

14. Benoit, L. et al. Astrocytes functionally integrate multiple synapses via specialized leaflet domains. Cell 188, 6453–6472.e16 (2025).

15. Boulay, A.-C. et al. Translation in astrocyte distal processes sets molecular heterogeneity at the gliovascular interface. Cell Discov. 3, (2017).

16. Hösli, L. et al. Direct vascular contact is a hallmark of cerebral astrocytes. Cell Rep. 39, 110599 (2022).

17. Stogsdill, J. A. et al. Astrocytic neuroligins control astrocyte morphogenesis and synaptogenesis. Nature 551, 192–197 (2017).

18. Lanjakornsiripan, D. et al. Layer-specific morphological and molecular differences in neocortical astrocytes and their dependence on neuronal layers. Nat. Commun. 9, 1623 (2018).

19. Abdeladim, L. et al. Multicolor multiscale brain imaging with chromatic multiphoton serial microscopy. Nat. Commun. 10.1038/s41467-019-09552-9 doi:10.1038/s41467-019-09552-9.

20. Clavreul, S. et al. Cortical astrocytes develop in a plastic manner at both clonal and cellular levels. Nat. Commun. 10, (2019).

21. García-Marqués, J. & López-Mascaraque, L. Clonal Identity Determines Astrocyte Cortical Heterogeneity. Cereb. Cortex 23, 1463–1472 (2013).

22. Salmon, C. K. et al. Organizing principles of astrocytic nanoarchitecture in the mouse cerebral cortex. Curr. Biol. CB 33, 957–972.e5 (2023).

23. Endo, F. et al. Molecular basis of astrocyte diversity and morphology across the CNS in health and disease. Science 378, eadc9020 (2022).

24. Dumas, L. et al. In Utero Electroporation of Multiaddressable Genome-Integrating Color (MAGIC) Markers to Individualize Cortical Mouse Astrocytes. J. Vis. Exp. 10.3791/61110 (2020) doi:10.3791/61110.

25. Zeisel, A. et al. Cell types in the mouse cortex and hippocampus revealed by single-cell RNA-seq. Science 347, 1138–1142 (2015).

26. Zeisel, A. et al. Molecular Architecture of the Mouse Nervous System. Cell 174, 999–1014.e22 (2018).

27. Bayraktar, O. A. et al. Astrocyte layers in the mammalian cerebral cortex revealed by a single-cell in situ transcriptomic map. Nat. Neurosci. 23, 500–509 (2020).

28. Batiuk, M. Y. et al. Identification of region-specific astrocyte subtypes at single cell resolution. Nat. Commun. 11, 1220 (2020).

29. Zhou, J. et al. Dual lineage origins contribute to neocortical astrocyte diversity. Nat. Commun. 16, 6992 (2025).

30. Lee, J. H. et al. Astrocyte morphogenesis requires self-recognition. Nature 644, 164–172 (2025).

31. Ge, W.-P., Miyawaki, A., Gage, F. H., Jan, Y. N. & Jan, L. Y. Local generation of glia is a major astrocyte source in postnatal cortex. Nature 484, 376–380 (2012).

32. Tsai, H.-H. et al. Regional Astrocyte Allocation Regulates CNS Synaptogenesis and Repair. Science 337, 358–362 (2012).

33. Anderson, S. A., Eisenstat, D. D., Shi, L. & Rubenstein, J. L. R. Interneuron Migration from Basal Forebrain to Neocortex: Dependence on Dlx Genes. Science 278, 474–476 (1997).

34. Anderson, S. A., Marín, O., Horn, C., Jennings, K. & Rubenstein, J. L. Distinct cortical migrations from the medial and lateral ganglionic eminences. Dev. Camb. Engl. 128, 353–363 (2001).

35. Marshall, C. A. G. & Goldman, J. E. Subpallial *Dlx2* -Expressing Cells Give Rise to Astrocytes and Oligodendrocytes in the Cerebral Cortex and White Matter. J. Neurosci. 22, 9821–9830 (2002).

36. Lozano Casasbuenas, D., et al. The laminar position, morphology, and gene expression profiles of cortical astrocytes are influenced by time of birth from ventricular/subventricular progenitors. Glia 72, 1693–1706 (2024).

37. Ojalvo-Sanz, A. C., Pernia-Solanilla, C. & López-Mascaraque, L. Spatial organization of astrocyte clones: The role of developmental progenitor timing. Glia 72, 1290–1303 (2024).

38. Mahou, P. et al. Multicolor two-photon tissue imaging by wavelength mixing. Nat. Methods 9, 815–818 (2012).

39. Srinivasan, R. et al. New Transgenic Mouse Lines for Selectively Targeting Astrocytes and Studying Calcium Signals in Astrocyte Processes In Situ and In Vivo. Neuron 92, 1181–1195 (2016).

40. Loulier, K. et al. Multiplex Cell and Lineage Tracking with Combinatorial Labels. Neuron 81, 505–520 (2014).

41. Kawaguchi, Y., Terashima, Y., Tanaka, S. & Okuda, H. Differential Expression Patterns of Astrocyte Markers in The Adult Mouse Brain. ACTA Histochem. Cytochem. 59, 123– 128 (2026).

42. Cahoy, J. D. et al. A Transcriptome Database for Astrocytes, Neurons, and Oligodendrocytes: A New Resource for Understanding Brain Development and Function. J. Neurosci. 28, 264–278 (2008).

43. Baldwin, K. T. et al. HepaCAM controls astrocyte self-organization and coupling. Neuron 109, 2427–2442.e10 (2021).

44. Kharlamova, A. et al. Spatial–temporal representation of the astroglial markers in the developing human cortex. Brain Struct. Funct. 229, 2385–2403 (2024).

45. Sun, W. et al. Sox9 Is an Astrocyte-Specific Nuclear Marker in the Adult Brain Outside the Neurogenic Regions. J. Neurosci. Off. J. Soc. Neurosci. 37, 4493–4507 (2017).

46. Susaki, E. A. et al. Advanced CUBIC protocols for whole-brain and whole-body clearing and imaging. Nat. Protoc. 10, 1709–1727 (2015).

47. Nery, S., Fishell, G. & Corbin, J. G. The caudal ganglionic eminence is a source of distinct cortical and subcortical cell populations. Nat. Neurosci. 5, 1279–1287 (2002).

48. Dumas, L., Durand, J. & Loulier, K. Multicolor Cell Lineage Tracing Using MAGIC Markers Strategies. Methods Mol. Biol. Clifton NJ 2886, 47–63 (2025).

49. Gilbert, A., Vidal, X. E., Estevez, R., Cohen-Salmon, M. & Boulay, A.-C. Postnatal development of the astrocyte perivascular MLC1/GlialCAM complex defines a temporal window for the gliovascular unit maturation. Brain Struct. Funct. 224, 1267–1278 (2019).

50. Gao, P. et al. Deterministic Progenitor Behavior and Unitary Production of Neurons in the Neocortex. Cell 159, 775–788 (2014).

51. Doyle, J. P. et al. Application of a Translational Profiling Approach for the Comparative Analysis of CNS Cell Types. Cell 135, 749–762 (2008).

52. Kim, E. J., Ables, J. L., Dickel, L. K., Eisch, A. J. & Johnson, J. E. Ascl1 (Mash1) Defines Cells with Long-Term Neurogenic Potential in Subgranular and Subventricular Zones in Adult Mouse Brain. PLoS ONE 6, e18472 (2011).

53. Marin, O., Anderson, S. A. & Rubenstein, J. L. Origin and molecular specification of striatal interneurons. J. Neurosci. Off. J. Soc. Neurosci. 20, 6063–6076 (2000).

54. Kim, E. J., Battiste, J., Nakagawa, Y. & Johnson, J. E. Ascl1 (Mash1) lineage cells contribute to discrete cell populations in CNS architecture. Mol. Cell. Neurosci. 38, 595– 606 (2008).

55. Liu, Y.-H. et al. Ascl1 promotes tangential migration and confines migratory routes by induction of Ephb2 in the telencephalon. Sci. Rep. 7, 42895 (2017).

56. Bandler, R. C. et al. Single-cell delineation of lineage and genetic identity in the mouse brain. Nature 601, 404–409 (2022).

57. Clavreul, S., Dumas, L. & Loulier, K. Astrocyte development in the cerebral cortex: Complexity of their origin, genesis, and maturation. Front. Neurosci. 16, 916055 (2022).

58. Magavi, S., Friedmann, D., Banks, G., Stolfi, A. & Lois, C. Coincident Generation of Pyramidal Neurons and Protoplasmic Astrocytes in Neocortical Columns. J. Neurosci. 32, 4762–4772 (2012).

59. Fogarty, M., Richardson, W. D. & Kessaris, N. A subset of oligodendrocytes generated from radial glia in the dorsal spinal cord. Development 132, 1951–1959 (2005).

60. García-Moreno, F. & Molnár, Z. Variations of telencephalic development that paved the way for neocortical evolution. Prog. Neurobiol. 194, 101865 (2020).

61. Bishop, K. M., Rubenstein, J. L. R. & O’Leary, D. D. M. Distinct Actions of *Emx1* , *Emx2* , and *Pax6* in Regulating the Specification of Areas in the Developing Neocortex. J. Neurosci. 22, 7627–7638 (2002).

62. Rueda-Alaña, E., Martínez-Garay, I., Encinas, J. M., Molnár, Z. & García-Moreno, F. Dbx1-Derived Pyramidal Neurons Are Generated Locally in the Developing Murine Neocortex. Front. Neurosci. 12, 792 (2018).

63. Tosches, M. A. et al. Evolution of pallium, hippocampus, and cortical cell types revealed by single-cell transcriptomics in reptiles. Science 360, 881–888 (2018).

64. Zaremba, B. et al. Developmental origins and evolution of pallial cell types and structures in birds. Science 387, eadp5182 (2025).

65. Majo, M., Koontz, M., Rowitch, D. & Ullian, E. M. An update on human astrocytes and their role in development and disease. Glia 68, 685–704 (2020).

66. deAzevedo, L. C., et al. Cortical radial glial cells in human fetuses: depth-correlated transformation into astrocytes. J. Neurobiol. 55, 288–298 (2003).

67. Bushong, E. A., Martone, M. E. & Ellisman, M. H. Maturation of astrocyte morphology and the establishment of astrocyte domains during postnatal hippocampal development. Int. J. Dev. Neurosci. 22, 73–86 (2004).

68. Heinrich, C. et al. Directing Astroglia from the Cerebral Cortex into Subtype Specific Functional Neurons. PLoS Biol. 8, e1000373 (2010).

69. Modo, M. Bioscaffold-Induced Brain Tissue Regeneration. Front. Neurosci. 13, 1156 (2019).

