## Supplementary Figures for "Neocortical astrocyte diversity stems from distinct developmental origins"

### Supplementary Fig. 1

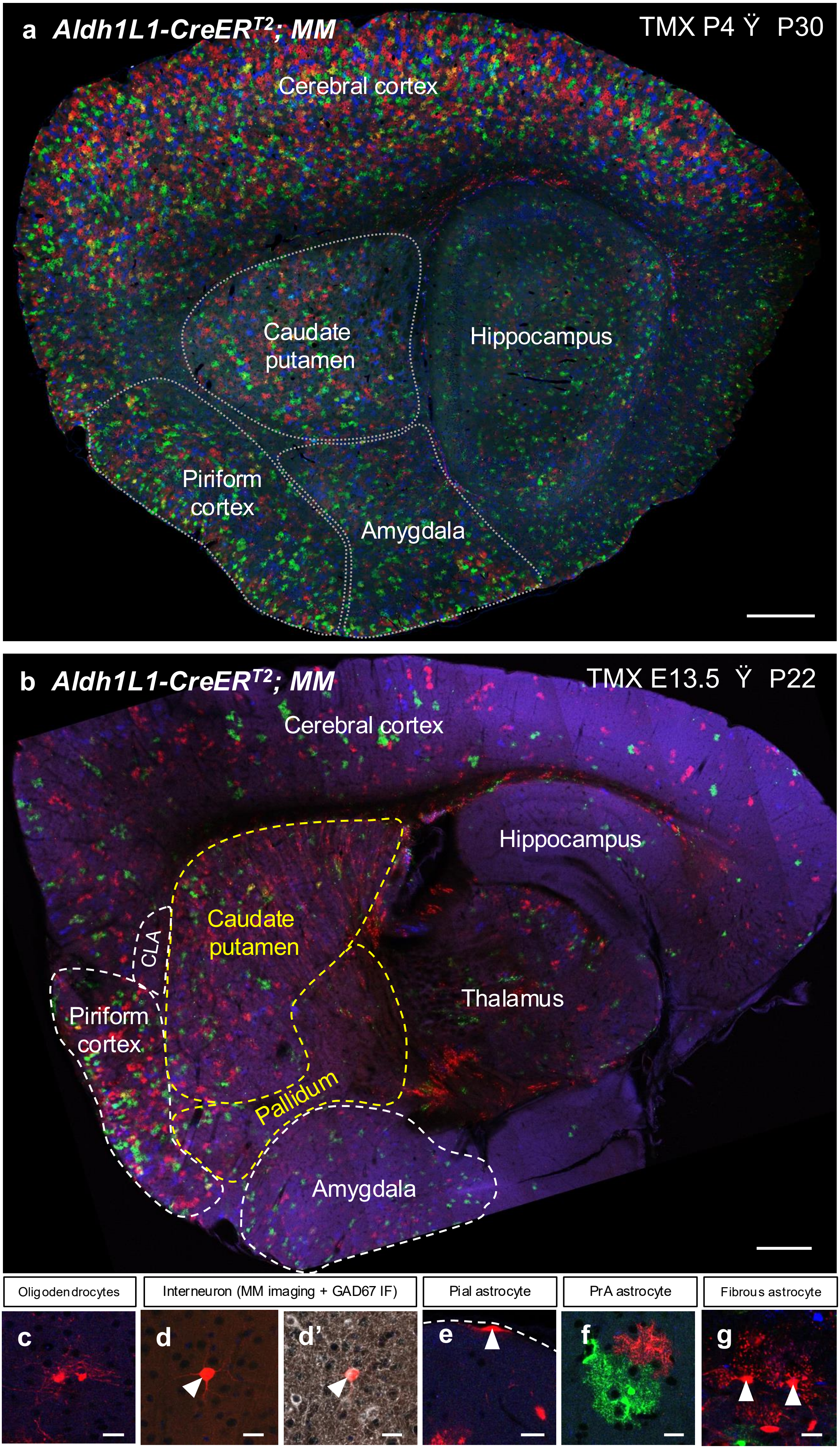

#### Supplementary Fig. 2

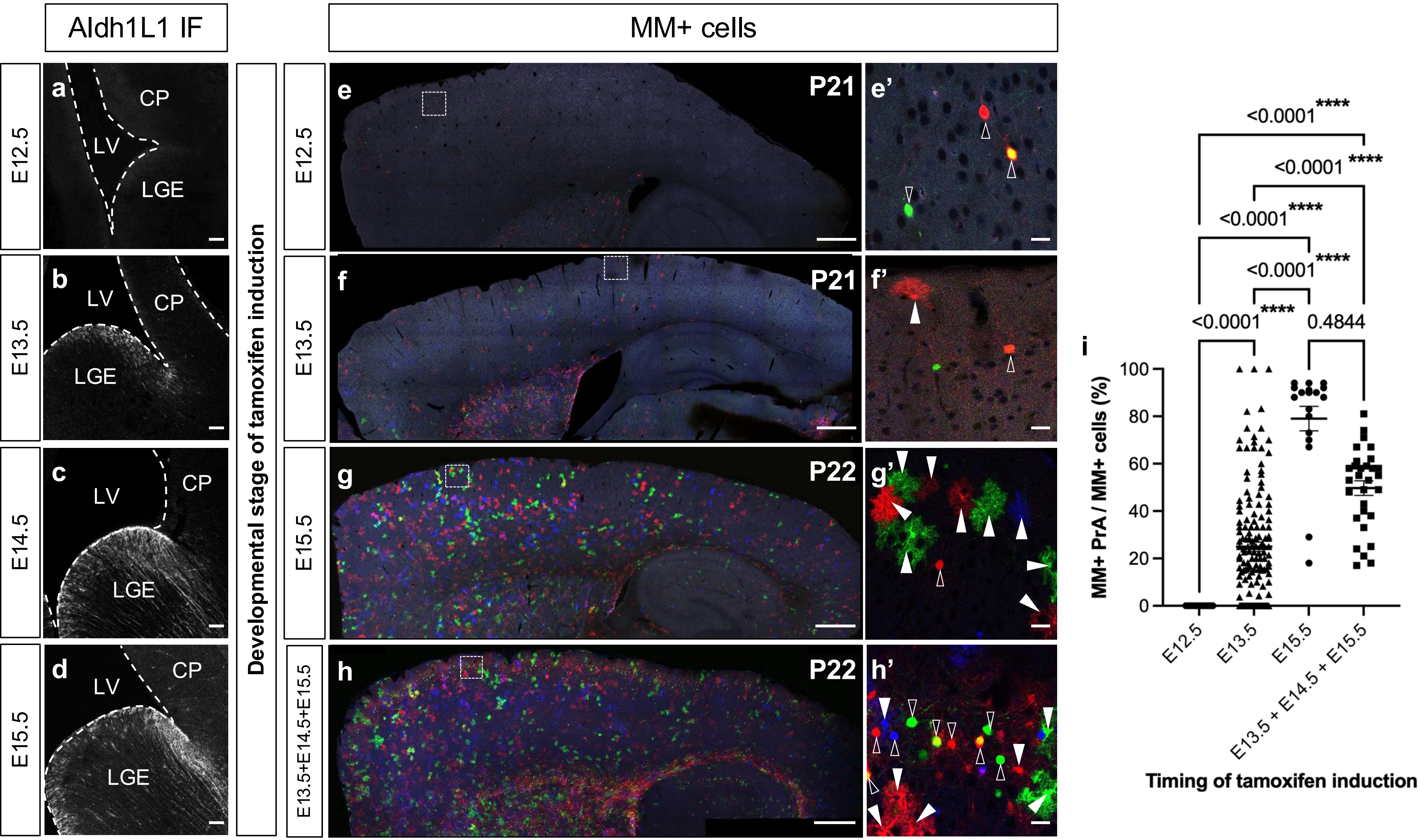

### Supplementary Fig. 3

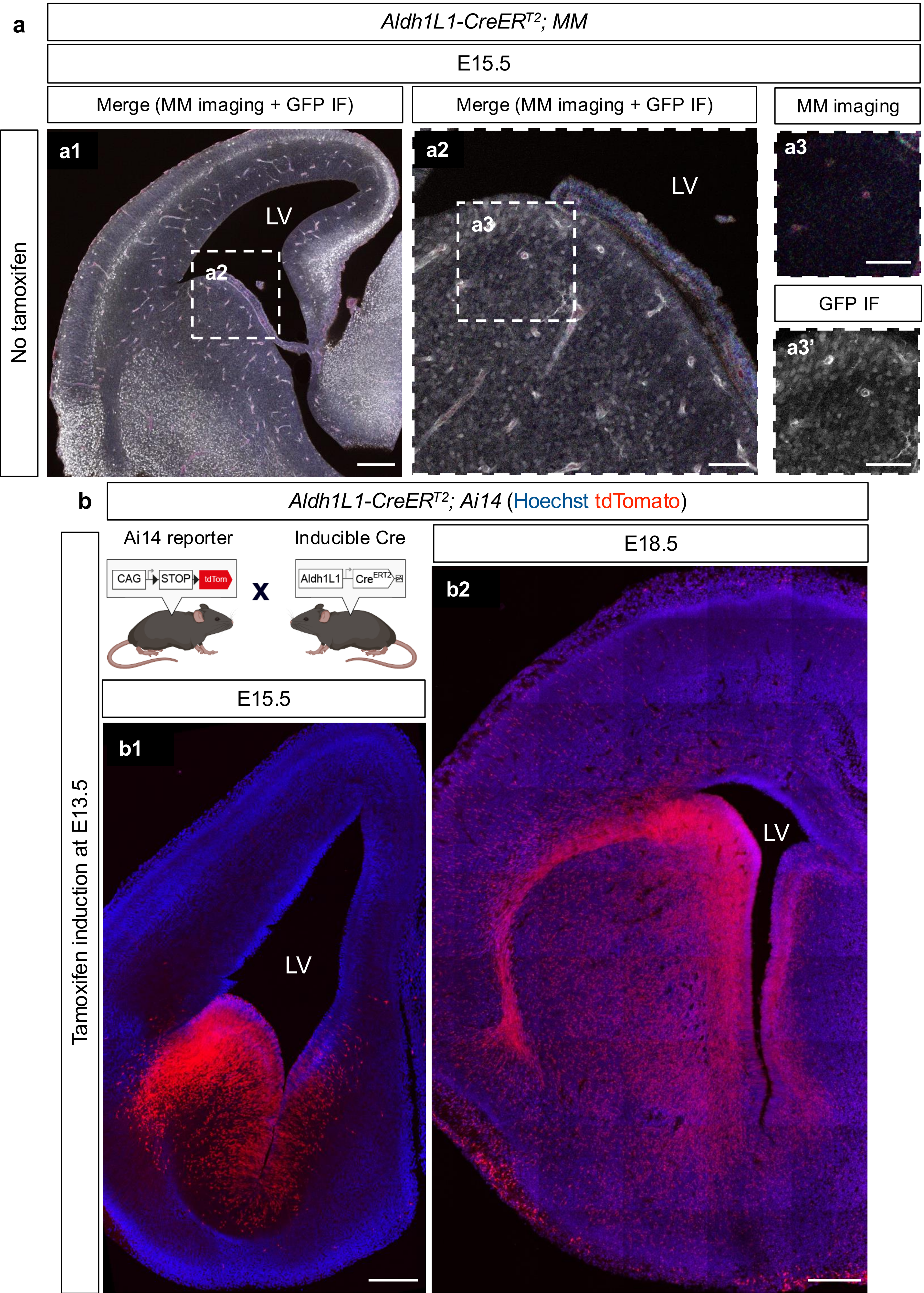

### *Supplementary Fig. 4*

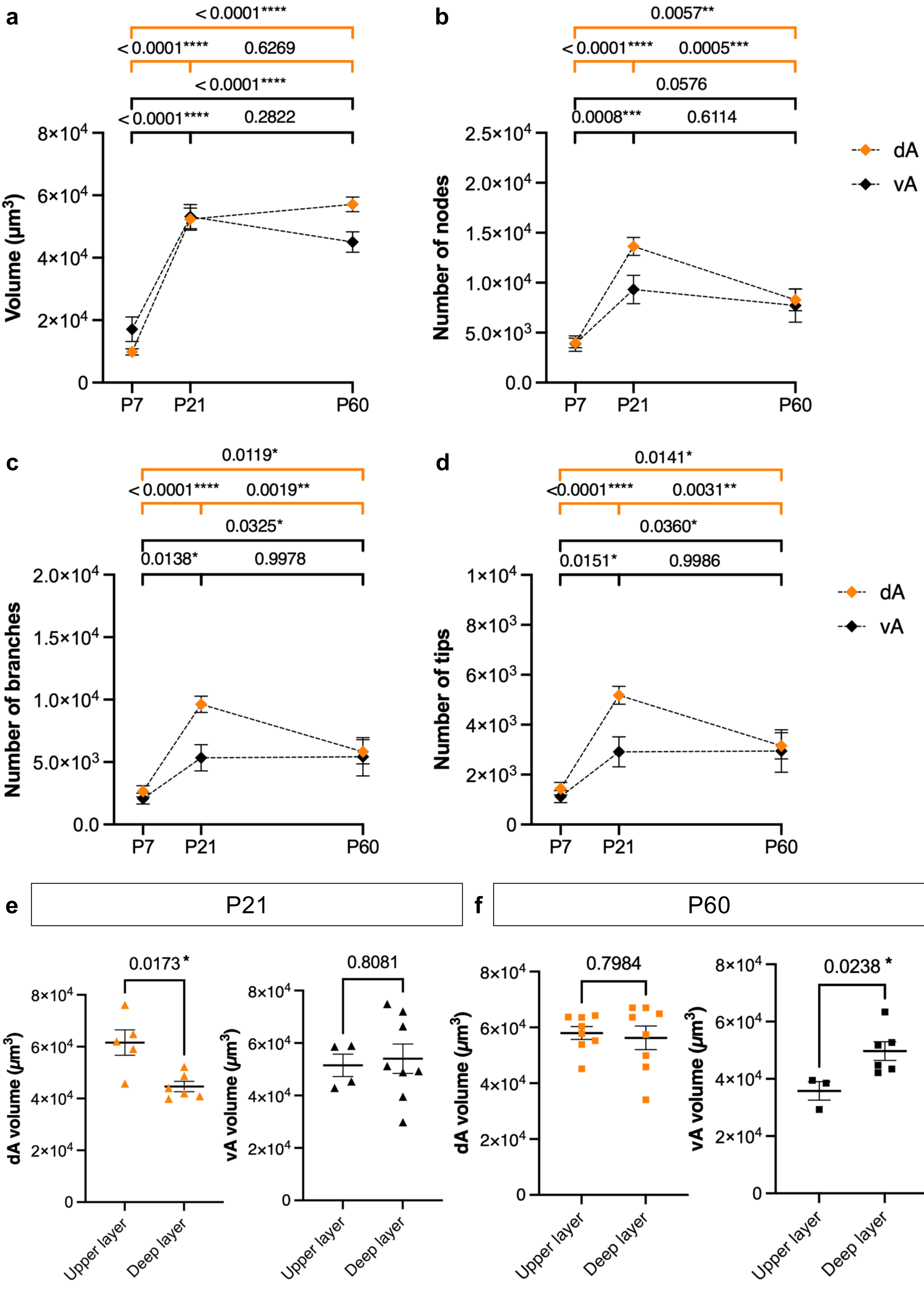

### Supplementary Fig. 5

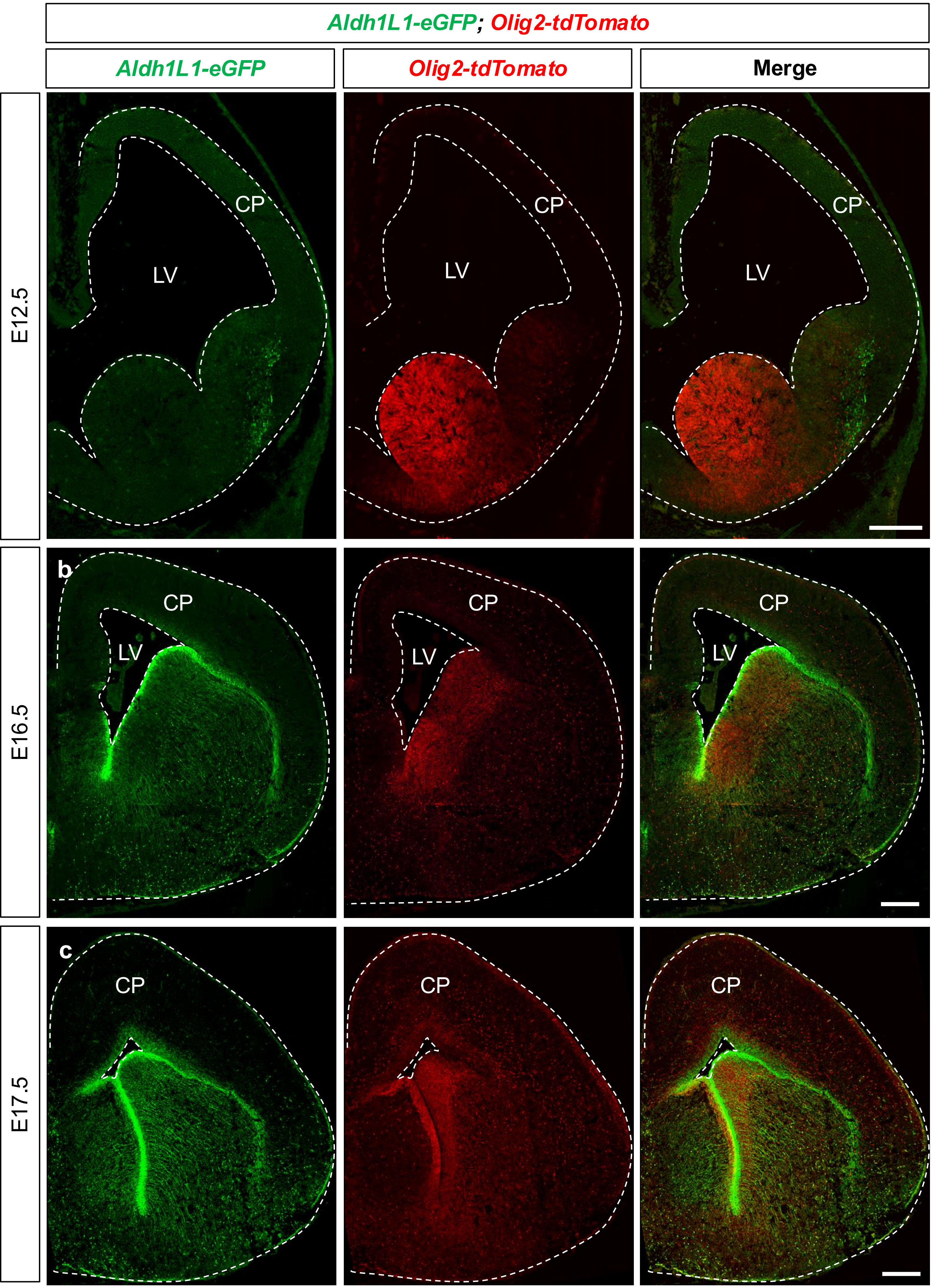

### Supplementary Fig. 6

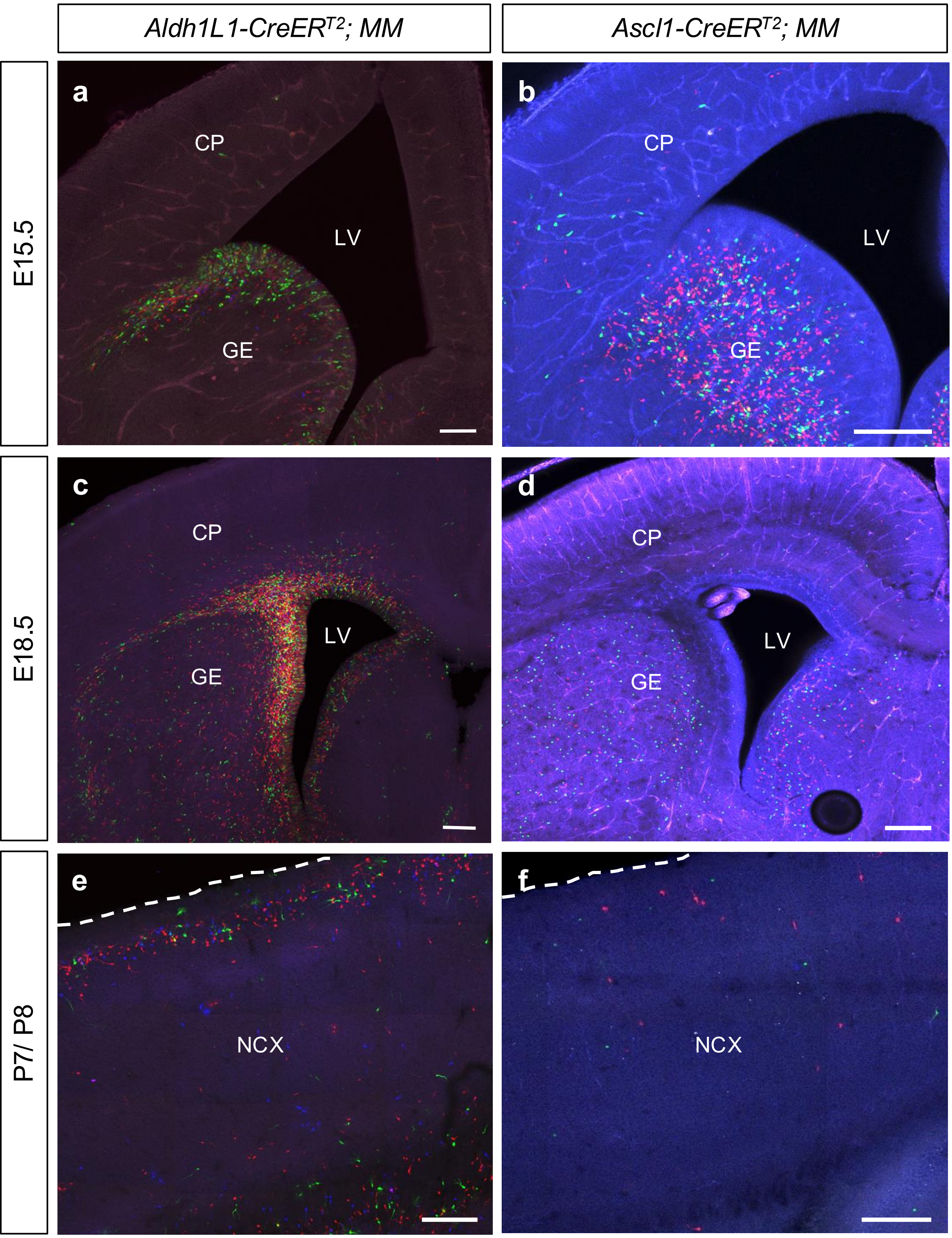
