## Supplementary Table for "Neocortical astrocyte diversity stems from distinct developmental origins"

| Major cell type | Gene |
|---|---|
| Ependymal cell | *Ccdc153* |
| Ependymal cell | *Tmem212* |
| Ependymal cell | *Dynlrb2* |
| Ependymal cell | *Pifo* |
| Ependymal cell | *Foxj1* |
| Astrocyte | *Aqp4* |
| Astrocyte | *Apoe* |
| Astrocyte | *Sox9* |
| Astrocyte | *Nfia* |
| Astrocyte | *Nfib* |
| Astrocyte | *S100b* |
| Astrocyte | *Aldh1l1* |
| Astrocyte | *Acsbg1* |
| Astrocyte | *Aldoc* |
| Astrocyte | *Atp1b2* |
| Astrocyte | *Fgfr3* |
| Astrocyte | *Slc1a3* |
| Astrocyte | *Gja1* |
| Neuron | *Rbfox3* |
| Neuron | *Mapt* |
| Neuron | *Syp* |
| Neuron | *Snap25* |
| Neuron | *Syt1* |
| Neuron | *Vamp2* |
| Neuron | *Rab3a* |
| Neuron | *Syn1* |
| Neuron | *Sv2a* |
| Neuron | *Sv2b* |
| Neuron | *Scamp5* |
| Neuron | *Dnm1* |
| Neuron | *Syngr1* |
| Neuron | *Syt4* |
| Neuron | *Snca* |
| Neuron | *Cck* |
| Neuron | *Map2* |
| Neuron | *Stmn2* |
| Neuron | *Syn1* |
| Microglia | *Ccl4* |
| Microglia | *Tnf* |
| Microglia | *Tmem119* |
| Microglia | *P2ry12* |
| Microglia | *Aif1* |
| Microglia | *Cd68* |
| Microglia | *Itgam* |
| Microglia | *Ptprc* |

| | |
|---|---|
| Microglia | *Cd86* |
| Microglia | *Trem2* |
| Microglia | *Cx3cr1* |
| Microglia | *Mafb* |
| Microglia | *Cd14* |
| Microglia | *Cd68* |
| Microglia | *Fcgr1* |
| Microglia | *Itgam* |
| Microglia | *Mertk* |
| Microglia | *Tlr2* |
| Pericyte | *Pdgfrb* |
| Pericyte | *Anpep* |
| Pericyte | *Des* |
| Pericyte | *Higd1b* |
| Pericyte | *Ndufa4l2* |
| Pericyte | *Rgs5* |
| Pericyte | *Cspg4* |
| Pericyte | *Mcam* |
| Endothelial cell | *Pecam1* |
| Endothelial cell | *Cd34* |
| Endothelial cell | *Tek* |
| Endothelial cell | *Kdr* |
| Endothelial cell | *Sema3g* |
| Endothelial cell | *Ly6c1* |
| Oligodendrocyte/OPC | *Sox10* |
| Oligodendrocyte/OPC | *Pdgfra* |
| Oligodendrocyte/OPC | *Lhfpl3* |
| Oligodendrocyte/OPC | *Gpr17* |
| Oligodendrocyte/OPC | *Tnr* |
| Oligodendrocyte/OPC | *Neu4* |
| Oligodendrocyte/OPC | *Tnr* |
| Oligodendrocyte/OPC | *Ccp110* |
| Oligodendrocyte/OPC | *Ninj2* |
| Oligodendrocyte/OPC | *Hapln2* |
| Oligodendrocyte/OPC | *Olig2* |
| Oligodendrocyte/OPC | *Mbp* |
| Oligodendrocyte/OPC | *Mog* |
| Oligodendrocyte/OPC | *Plp1* |
| Oligodendrocyte/OPC | *Mag* |
| Oligodendrocyte/OPC | *Olig1* |
| Radial glial cell | *Pclaf* |
| Radial glial cell | *Cenpf* |
| Radial glial cell | *Hells* |
| Radial glial cell | *Fabp7* |
| Radial glial cell | *Notch1* |
| Radial glial cell | *Top2a* |
| Radial glial cell | *Mki67* |
| Radial glial cell | *Ube2c* |

| | |
|---|---|
| Radial glial cell | *Ascl1* |
| Radial glial cell | *Cdk6* |
| Radial glial cell | *Mcm2* |
| Radial glial cell | *Egfr* |
| Radial glial cell | *Uhrf2* |
| Neuroblast | *Dcx* |
| Neuroblast | *Stmn2* |
| Neuroblast | *Celf4* |
| Neuroblast | *Islr2* |
| Neuroblast | *Tubb3* |
| Neuroblast | *Igfbpl1* |
| Neuroblast | *Tiam2* |
| Neuroblast | *Cd24a* |
| Neuroblast | *Dlx2* |
| Neuroblast | *Fxyd6* |
| Neuroblast | *Ncam1* |
| Proliferating cell | *Anln* |
| Proliferating cell | *Anp32e* |
| Proliferating cell | *Apc* |
| Proliferating cell | *Atad2* |
| Proliferating cell | *Aurka* |
| Proliferating cell | *Birc5* |
| Proliferating cell | *Blm* |
| Proliferating cell | *Casp8ap2* |
| Proliferating cell | *Cbx5* |
| Proliferating cell | *Ccna2* |
| Proliferating cell | *Ccnb1* |
| Proliferating cell | *Ccnb2* |
| Proliferating cell | *Ccnd2* |
| Proliferating cell | *Ccne1* |
| Proliferating cell | *Ccne2* |
| Proliferating cell | *Ccni* |
| Proliferating cell | *Ccnt1* |
| Proliferating cell | *Ccnt2* |
| Proliferating cell | *Ccny* |
| Proliferating cell | *Cdc20* |
| Proliferating cell | *Cdc25c* |
| Proliferating cell | *Cdc45* |
| Proliferating cell | *Cdca2* |
| Proliferating cell | *Cdca3* |
| Proliferating cell | *Cdca7* |
| Proliferating cell | *Cdca8* |
| Proliferating cell | *Cdk1* |
| Proliferating cell | *Cdk16* |
| Proliferating cell | *Cdk2* |
| Proliferating cell | *Cdk4* |
| Proliferating cell | *Cdk7* |
| Proliferating cell | *Cenpa* |

| | |
|---|---|
| Proliferating cell | *Cenpe* |
| Proliferating cell | *Cenpf* |
| Proliferating cell | *Chaf1b* |
| Proliferating cell | *Ckap2* |
| Proliferating cell | *Ckap2l* |
| Proliferating cell | *Ckap5* |
| Proliferating cell | *Cks1b* |
| Proliferating cell | *Cks2* |
| Proliferating cell | *Clspn* |
| Proliferating cell | *Ctcf* |
| Proliferating cell | *Dlgap5* |
| Proliferating cell | *Dtl* |
| Proliferating cell | *Ect2* |
| Proliferating cell | *Fen1* |
| Proliferating cell | *G2e3* |
| Proliferating cell | *Gas2l3* |
| Proliferating cell | *Gins2* |
| Proliferating cell | *Gmnn* |
| Proliferating cell | *Gtse1* |
| Proliferating cell | *Hells* |
| Proliferating cell | *Hjurp* |
| Proliferating cell | *Hmgb2* |
| Proliferating cell | *Hmmr* |
| Proliferating cell | *Hn1* |
| Proliferating cell | *Kif11* |
| Proliferating cell | *Kif20b* |
| Proliferating cell | *Kif23* |
| Proliferating cell | *Kif2c* |
| Proliferating cell | *Lbr* |
| Proliferating cell | *Mcm2* |
| Proliferating cell | *Mcm4* |
| Proliferating cell | *Mcm5* |
| Proliferating cell | *Mcm6* |
| Proliferating cell | *Mki67* |
| Proliferating cell | *Msh2* |
| Proliferating cell | *Nasp* |
| Proliferating cell | *Ncapd2* |
| Proliferating cell | *Nek2* |
| Proliferating cell | *Nuf2* |
| Proliferating cell | *Nusap1* |
| Proliferating cell | *Pcna* |
| Proliferating cell | *Pola1* |
| Proliferating cell | *Pold3* |
| Proliferating cell | *Prim1* |
| Proliferating cell | *Psrc1* |
| Proliferating cell | *Rad51* |
| Proliferating cell | *Rad51ap1* |
| Proliferating cell | *Rangap1* |

| | |
|---|---|
| Proliferating cell | *Rfc2* |
| Proliferating cell | *Rpa2* |
| Proliferating cell | *Rrm1* |
| Proliferating cell | *Rrm2* |
| Proliferating cell | *Slbp* |
| Proliferating cell | *Smc4* |
| Proliferating cell | *Tacc3* |
| Proliferating cell | *Tipin* |
| Proliferating cell | *Tmpo* |
| Proliferating cell | *Top2a* |
| Proliferating cell | *Tpx2* |
| Proliferating cell | *Tubb4b* |
| Proliferating cell | *Tyms* |
| Proliferating cell | *Ube2c* |
| Proliferating cell | *Ubr7* |
| Proliferating cell | *Uhrf1* |
| Proliferating cell | *Ung* |
| Proliferating cell | *Usp1* |
| Proliferating cell | *Wdr76* |
